# Impact of the B.1.1.7 variant on neutralizing monoclonal antibodies recognizing diverse epitopes on SARS-CoV-2 Spike

**DOI:** 10.1101/2021.02.03.429355

**Authors:** Carl Graham, Jeffrey Seow, Isabella Huettner, Hataf Khan, Neophytos Kouphou, Sam Acors, Helena Winstone, Suzanne Pickering, Rui Pedro Galao, Maria Jose Lista, Jose M Jimenez-Guardeno, Adam G. Laing, Yin Wu, Magdalene Joseph, Luke Muir, Weng M. Ng, Helen M. E. Duyvesteyn, Yuguang Zhao, Thomas A. Bowden, Manu Shankar-Hari, Annachiara Rosa, Peter Cherepanov, Laura E. McCoy, Adrian C. Hayday, Stuart J.D. Neil, Michael H. Malim, Katie J. Doores

## Abstract

The interaction of the SARS-CoV-2 Spike receptor binding domain (RBD) with the ACE2 receptor on host cells is essential for viral entry. RBD is the dominant target for neutralizing antibodies and several neutralizing epitopes on RBD have been molecularly characterized. Analysis of circulating SARS-CoV-2 variants has revealed mutations arising in the RBD, the N-terminal domain (NTD) and S2 subunits of Spike. To fully understand how these mutations affect the antigenicity of Spike, we have isolated and characterized neutralizing antibodies targeting epitopes beyond the already identified RBD epitopes. Using recombinant Spike as a sorting bait, we isolated >100 Spike-reactive monoclonal antibodies from SARS-CoV-2 infected individuals. ~45% showed neutralizing activity of which ~20% were NTD-specific. None of the S2-specific antibodies showed neutralizing activity. Competition ELISA revealed that NTD-specific mAbs formed two distinct groups: the first group was highly potent against infectious virus, whereas the second was less potent and displayed glycan-dependant neutralization activity. Importantly, mutations present in B.1.1.7 Spike frequently conferred resistance to neutralization by the NTD-specific neutralizing antibodies. This work demonstrates that neutralizing antibodies targeting subdominant epitopes need to be considered when investigating antigenic drift in emerging variants.

## Introduction

Severe acute respiratory syndrome coronavirus 2 (SARS-CoV-2) is the causative agent of COVID-19. SARS-CoV-2 belongs to the *Betacoronavirus* genus of the *Coronaviridae* family, alongside severe acute respiratory syndrome coronavirus (SARS-CoV) and Middle East respiratory syndrome coronavirus (MERS-CoV) (Lu et al., 2020). The positive sense RNA genome encodes four structural proteins; Spike (S), envelope (E), membrane (M) and nucleocapsid (N) (Jiang et al., 2020). The S glycoprotein is responsible for interaction with the human angiotensin-converting enzyme 2 (ACE2) receptor and subsequent virus-cell membrane fusion and thus is the key target for neutralizing antibodies (Pinto et al., 2020). The Spike glycoprotein assembles into homotrimers on the viral membrane, with each Spike monomer encompassing two functional subunits, S1 and S2. The S1 subunit contains the N-terminal domain (NTD) and the receptor binding domain (RBD). The RBD encompasses the receptor binding motif (RBM) that directly contacts the ACE2 receptor. The S2 subunit, containing the fusion peptide, two heptad repeats (HR1 and HR2), the cytoplasmic tail and the transmembrane domain, is crucial for viral membrane fusion (Yao et al., 2020).

Despite the recent emergence of SARS-CoV-2 in the human population, a rapid understanding of the antibody response arising from infection has emerged (Beaudoin-Bussieres et al., 2020; Crawford et al., 2020; Dan et al., 2021; Muecksch et al., 2020; Okba et al., 2020; Pickering et al., 2020; Prevost et al., 2020; Seow et al., 2020). The majority of SARS-CoV-2 infected individuals have been shown to generate an antibody response 5-15 days post onset of symptoms (POS) that peaks after ~3-4 weeks and then starts to decline (Crawford et al., 2020; Muecksch et al., 2020; Prevost et al., 2020; Seow et al., 2020). The magnitude of the neutralizing antibody response, which is thought to be important for protection from re-infection and/or disease, has been associated with disease severity. Specifically, those with most severe disease typically develop the strongest antibody response whereas those experiencing mild disease, or who are asymptomatic, can have lower levels of neutralizing activity detectable in their sera (Guthmiller et al., 2021; Laing et al., 2020b; Legros et al., 2021; Rees-Spear et al., 2021; Seow et al., 2020; Zohar et al., 2020). Antibodies targeting RBD have been suggested to account for >90% of neutralizing activity in convalescent sera (Greaney et al., 2020; Piccoli et al., 2020) and several neutralizing epitopes on RBD that are targeted by highly potent monoclonal antibodies have been molecularly characterized (Barnes et al., 2020; Brouwer et al., 2020; Piccoli et al., 2020; Pinto et al., 2020; Rogers et al., 2020; Tortorici et al., 2020; Wang et al., 2020; Wu et al., 2020). Reports suggest escape from RBD-mediated neutralization is occurring in variant strains that are emerging globally, which include mutations within the RBD that have been postulated to enable escape (Muecksch et al., 2020; Starr et al., 2021; Wimber et al., 2021). This highlights the need to identify neutralizing antibodies that bind epitopes outside RBD and to understand the role these antibodies play in protection from re-infection or following vaccination. Identification of neutralizing epitopes beyond RBD is therefore important for the development of synergistic antibody cocktails for immunotherapy and passive vaccination, and will also be critical for evaluating the relevance of potential immune escape viral variants as they arise, for example the recently identified B.1.1.7 (Rambaut et al., 2020). We therefore sought to isolate SARS-CoV-2 neutralizing antibodies from three donors experiencing severe, mild or asymptomatic COVID-19 using an un-cleaved, pre-fusion stabilized trimeric Spike glycoprotein as antigen-bait to further characterize the neutralizing epitopes present on SARS-CoV-2 Spike.

Here, we isolated 107 mAbs across three donors of which 47 (43.9%) showed neutralizing activity. The majority (72.3%, 34/47) of the neutralizing antibodies targeted the RBD and formed four distinct competition groups. 21.3% (10/47) of neutralizing antibodies targeted the NTD and formed two separate groups. One NTD group contained potent neutralizing antibodies able to neutralize infectious virus at a similar potency to RBD-targeted neutralizing antibodies. The second NTD group, although less potent, showed glycan-dependant neutralization activity. NTD-specific neutralizing antibodies (nAbs) showed a dramatic decrease in neutralization potency against the recently reported highly transmissible B.1.1.7 variant, whereas RBD-specific nAbs were either unaffected or showed lower decreases in potency, indicating that neutralizing antibodies against epitopes outside RBD are important to consider when investigating antigenic drift surveillance and identifying newly emerging SARS-CoV-2 variants of concern.

## Results

### Ab responses to SARS-CoV-2 infection in three donors with varied COVID-19 symptoms

We have previously studied antibody responses following SARS-CoV-2 infection (Laing et al., 2020a; Seow et al., 2020). To compare the antibody response between different disease severities at the monoclonal level, we selected three donors experiencing a range of COVID-19 severity (Characterisation and Management of, 2020). P003 was hospitalized and spent time in ICU, P054 was symptomatic but did not require hospitalization and P008 was asymptomatic and SARS-CoV-2 infection was only identified through serology screening (Laing et al., 2020a). Longitudinal plasma samples were used to measure binding and neutralization titres (**Figure 1A**). As we and others have previously shown (Seow et al., 2020), the highest neutralization titre was detected in the individual with most severe disease (ID_50_ 9,181) and the lowest neutralization titre in the asymptomatic individual (ID_50_ 820). The nAb response declined during the convalescent period with neutralizing ID_50_ values reduced to 258 in P054 and 25 in P008 after 188- and 194-days post onset of symptoms respectively. Plasma IgG, IgM and IgA binding to Spike and RBD were also measured using ELISA, and although IgG to Spike and RBD remained detectable, a large decrease from peak binding was observed (**Figure 1A**).

**Figure 1:**
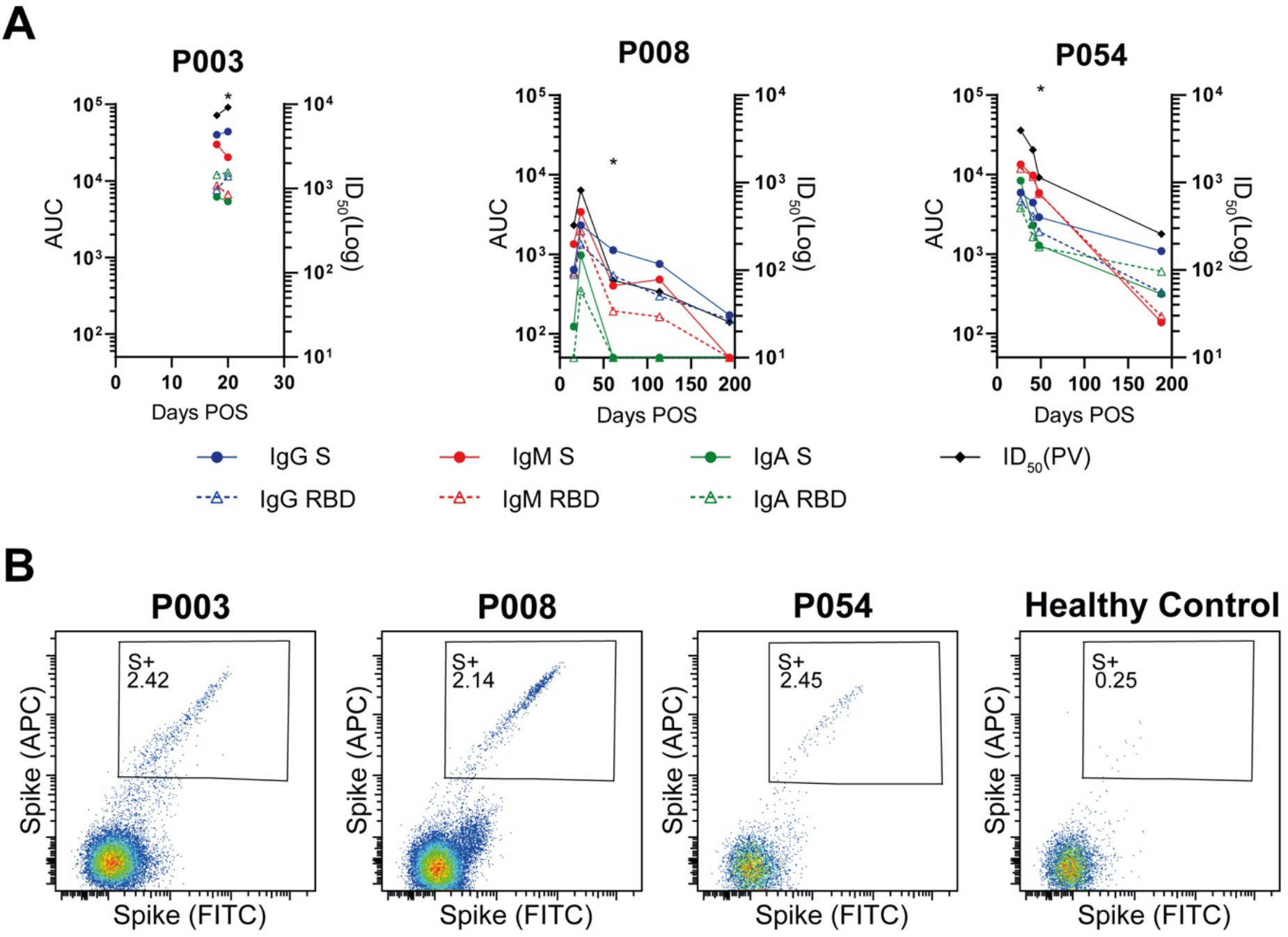
Binding and neutralization properties of plasma from donors used for B cell sorting and characterization of SARS-CoV-2 Spike reactive IgG+ B cells. **A)** Kinetics of the antibody binding response (IgM, IgA, IgG against S, RBD) and neutralization activity against SARS-CoV-2 pseudovirus for donors P003, P008 and P0054 in the acute and convalescent phase of SARS-CoV-2 infection. ELISA data is reported as area under the curve (AUC, left y-axis). Neutralization ID_50_ against pseudovirus is shown on the right x-axis. The asterix indicates the timepoint from which monoclonal antibodies were cloned for each donor. Experiments were performed in duplicate and repeated twice where plasma was available. **B)** Fluorescent activated cell sorting (FACS) showing percentage of CD19^+^IgG^+^ B Cells binding to SARS-CoV-2 Spike. The full gating strategy can be found in **Supplementary Figure S1**. A healthy control PBMC sample collected prior to the COVID-19 pandemic was used as a control to measure background binding to Spike.

### Antibodies generated following SARS-CoV-2 infection target RBD and other epitopes on S1 and S

Next, we used antigen-specific B cell sorting to isolate mAbs specific for SARS-CoV-2 Spike. PBMCs were available for sorting at days 20, 48, 61 POS from donors P003, P054 and P008, respectively. We used an uncleaved Spike that was stabilised in the prefusion conformation (GGGG substitution at furin cleavage site and 2P mutation (Wrapp et al., 2020)) as sorting bait to allow isolation and characterization of mAbs against the full range of neutralizing and non-neutralizing epitopes. 2.39%, 2.14%, and 2.45% of CD19+IgG+ B cells bound to Spike in donors P003, P008 and P054, respectively, compared to 0.25% for a pre-COVID-19 healthy control sample (**Figure 1B** and **Supplementary Figure 1**). Despite a low ID_50_ of 76 for P008 at day 61 POS, Spike reactive IgG^+^ B cells were detected at a similar frequency to P054 with an ID_50_ of 1,144. 150 Spike-reactive cells were sorted from donors P003 and P008 and 120 cells from donor P054. The heavy and light chains were reverse transcribed and amplified using nested PCR with gene specific primers. The purified PCR products were ligated into heavy and light chain expression vectors using Gibson assembly and the ligation products directly transfected into 293T cells (Rogers et al., 2020). Spike-reactive mAbs were identified by measuring Spike binding and IgG expression of supernatants using ELISA. The transformed Gibson products for Spike reactive mAbs were sequenced and used for gene analysis (see **Figure 3** and **Supplementary Figure 2**). Small scale expression of sequenced antibodies was used to determine specificity towards Spike, S1, NTD and RBD in ELISA (**Figure 2A**). In total, 107 Spike-reactive mAbs were identified and sequenced, 24, 19 and 64 from donors P003, P054 and P008 respectively (**Figure 2B**). 38/107 (35.5%) of the Spike reactive mAbs were RBD-specific, 35/107 (32.7%) were NTD-specific, and 1/107 (0.9%) bound S1 only (**Figure 2C**). 33/107 (30.8%) mAbs only bound Spike suggesting these mAbs are either specific for S2 or bind quaternary epitopes that span multiple subunits (Liu et al., 2020). The distribution of mAb epitopes targeted differed between donors, with P003 mAbs predominantly binding non-S1 epitopes (**Figure 2D**).

**Figure 2:**
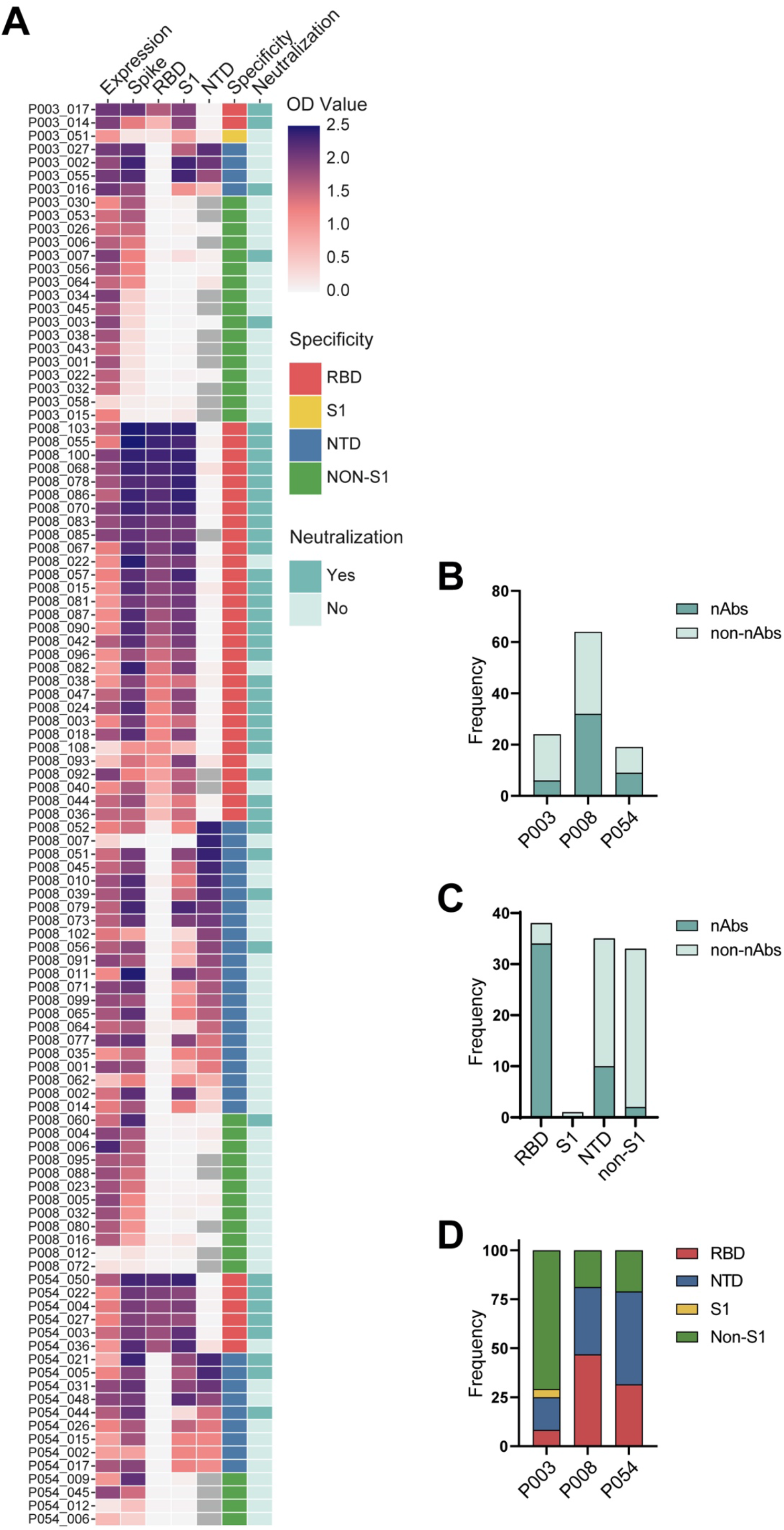
Identification of 107 SARS-CoV-2 Spike reactive monoclonal antibodies from three convalescent donors. **A)** Heat map showing IgG expression level and binding to SARS-CoV-2 Spike protein and the three subdomains S1, NTD and RBD. The figure reports OD values (range 0-2.5) for undiluted supernatant from small scale transfection of verified sequences of 107 cloned mAbs. Grey indicates binding not measured. Antigen binding was considered positive when OD at 405 nm >0.3 after background was subtracted. Neutralization activity was measured against pseudotyped virus using either small scale purified IgG or concentrated supernatant. Antibodies were considered neutralizing if at least 50% neutralization was reached at the highest concentration (5 μg/mL for purified mAb) or concentrated supernatant (30 times concentrated). Grey squares indicate samples that were not measured for the antigen. **B)** Bar graph showing frequency of neutralizing and nonneutralizing antibodies isolated from donors P003, P008 and P054. **C)** Bar graph showing frequency of neutralizing and non-neutralizing antibodies targeting specific sub-domains of Spike (RBD, NTD, S1, Non-S1). **D)** Bar graph showing the % of mAbs isolated from each donor targeting specific sub-domains of Spike (RBD, NTD, S1, Non-S1).

**Figure 3:**
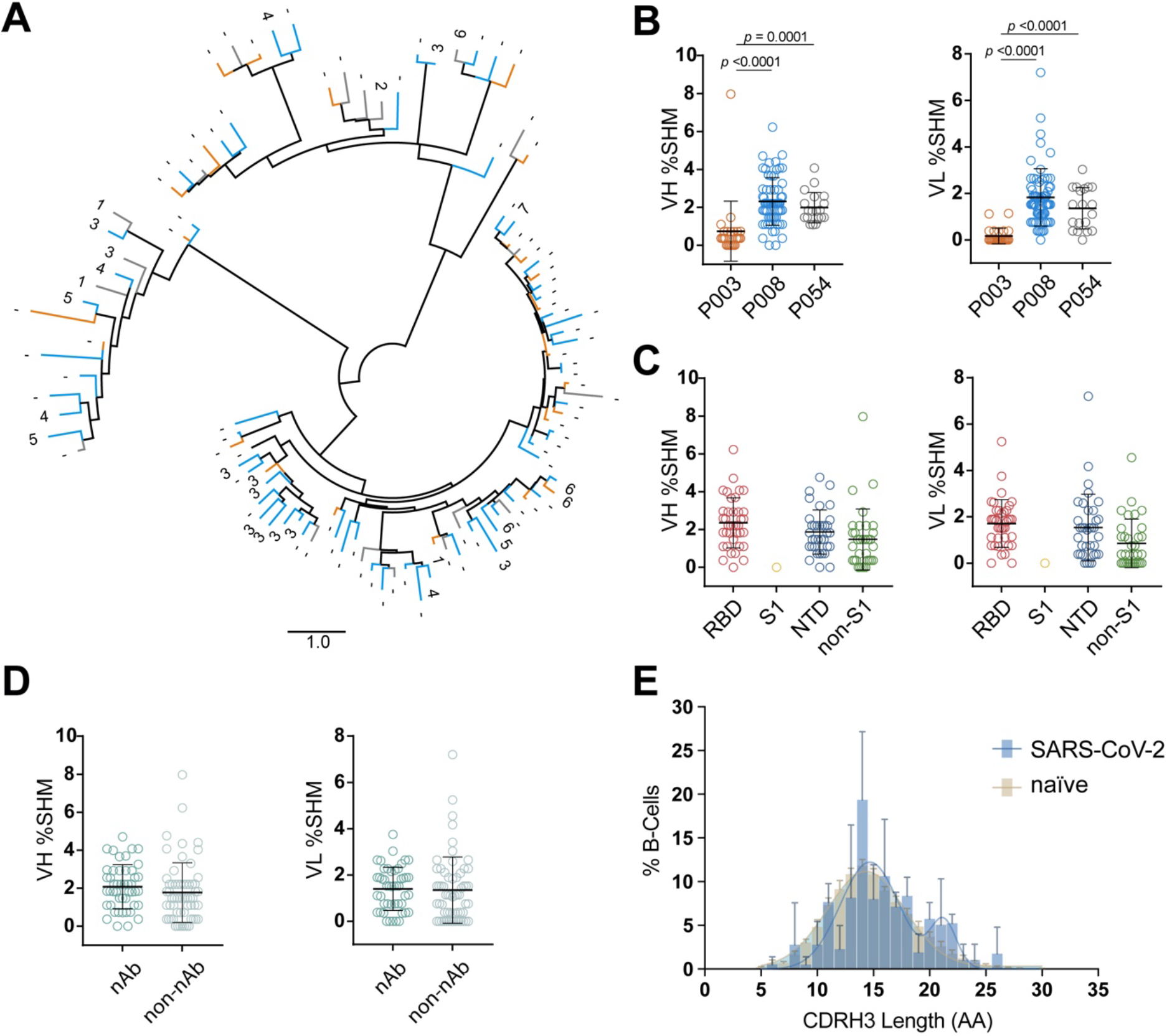
Sequence analysis of SARS-CoV-2 Spike specific monoclonal antibodies. **A)** Phylogenetic tree generated through analysis of heavy chain amino acid sequences. The first 5 amino acids were excluded in this analysis due to the variability introduced with the primer cocktail used during PCR. The sequences were aligned with Clustal W and clustered via PhyML to produce maximum likelihood phylogenetic trees which were visualized and annotated using FigTree. Branches are colour coded based on the donor each mAb was isolated from (P003 – orange, P008 – blue, P054 – grey). Binding competition groups (1-7) are labelled at the end of branches for those mAbs studies in competition analysis (**Figure 4A**). **B)** Level of somatic hypermutation (SHM) in the VH and VL genes for donors P003 (orange), P008 (blue) and P054 (grey). Differences between groups were determined using Kruskal-Wallis multiple comparison test and *p* values <0.05 shown. Black lines represent the mean SHM. **C)** Level of SHM for mAb targeting the RBD, NTD, S1 or non-S1 epitopes. No statistical difference in SHM was observed between the four groups for VH or VL (Kruskal-Wallis multiple comparison test). Black lines represent the mean SHM. **D)** Level of VH and VL SHM for neutralizing and non-neutralizing antibodies. No statistical difference was observed between the two groups (Mann-Whitney 2-sided U-test). Black lines represent the mean SHM. **E)** Distribution of CDRH3 lengths for SARS-CoV-2 specific mAbs and representative naïve B cell repertoire (Briney et al., 2019). Error bars represent the standard deviation between donors used in the analysis (n = 3 for SARS-CoV-2 and n=10 for naïve repertoire). A bimodal distribution of CDRH3 length is observed for SARS-CoV-2 Spike reactive mAbs.

SARS-CoV-2 neutralization activity of small scale purified mAb or concentrated supernatant was determined using an HIV-1 (human immunodeficiency virus type-1) based virus particles, pseudotyped with SARS-CoV-2 Spike (Grehan et al., 2015; Thompson, 2020) and a HeLa cell-line stably expressing the ACE2 receptor (Seow et al., 2020). 47/107 (43.9%) of cloned mAbs had neutralizing activity, an observation which highlights the presence of exposed but non-neutralizing epitopes on Spike, including RBD, that generate a strong antibody response. 34/37 (91.9%) of RBD-specific mAbs were neutralizing whereas only 10/35 (28.6%) of NTD-specific, and 3/34 (8.8%) of the Non-S1 binding mAbs had neutralizing activity (**Figure 2A and 2C**). Therefore, RBD was the dominant target for the neutralizing antibodies isolated in this study consistent with prior literature (Piccoli et al., 2020).

### Diversity in Ab gene usage

All Spike-reactive mAbs were sequenced and their heavy and light germline gene usage, level of somatic hypermutation (SHM) and CDRH3 length determined using the ImmunoGenetics (IMGT) database (Brochet et al., 2008). A diverse range of heavy and light chain germline genes were utilized by mAbs isolated from the three donors (**Supplementary Figure 2A**). Although an enrichment for certain germline genes was observed (**Supplemental Figure 2B**), there was only one example of clonal expansion arising from donor P008 (**Supplementary Figure 2D**). A comparison of VH gene usage compared to that of naïve B cell repertoires (Briney et al., 2019) showed an enrichment in VH3-30 usage and a de-enrichment in VH3-23 and to a lesser extent VH3-7 (**Supplementary Figure 2C**). Despite the enrichment in VH3-30 gene usage, ten different light chains gene pairings were observed, including both kappa and lambda genes (**Supplementary Figure 2B**). mAbs encoded by the VH 3-66 and 3-53 germlines were frequently observed for RBD-specific mAbs as previously described (Barnes et al., 2020; Kim et al., 2021; Robbiani et al., 2020; Yuan et al., 2020a). A phylogenetic tree of heavy chains showed clustering of related sequences from all three donors (**Figure 3A**).

Overall, low levels of SHM were observed in the VH and VL genes (mean of 1.9% and 1.4%, respectively) which is expected following an acute viral infection. There was statistically higher SHM in VH of mAbs from P008 (2.3%) and P054 (2.0%), which used PBMCs isolated at days 61 and 48 POS respectively, compared to P003 (0.8%) which used PBMC isolated at day 20 POS (**Figure 3B**). There was no difference in the level of SHM in the heavy or light chains between RBD-specific, NTD-specific and non-S1 mAbs (**Figure 3C**) or between neutralizing and non-neutralizing mAbs (**Figure 3D**). Comparison of the CDR3 length distribution with representative naïve repertoires showed an enrichment in CDRH3 of lengths 21 and 22 in the

SARS-CoV-2 specific mAbs (**Figure 3E**). Overall, and similar to previously reported (Barnes et al., 2020; Brouwer et al., 2020; Jiang et al., 2020; Liu et al., 2020; Pinto et al., 2020; Robbiani et al., 2020; Rogers et al., 2020; Tortorici et al., 2020; Wu et al., 2020), the repertoire of SARS-CoV-2 specific mAbs was very diverse, did not differ greatly from that observed in representative naïve repertoires and showed very little SHM.

### Neutralizing activity of SARS-CoV-2 mAbs

Twelve non-neutralizing and 37 neutralizing mAbs were selected for large scale expression/purification and further characterization. The neutralization potency was measured against both pseudoviral particles and infectious virus (PHE strain using Vero-E6 as target cells). IC_50_ values ranged from 1.2 - 660 ng/mL against the pseudovirus particles and 2.3 - 488 ng/mL against infectious virus (**Supplementary Table S1**).

RBD-specific mAb P008_108 is amongst the most potent anti-SARS-CoV-2 mAbs described so far with an IC_50_ of 2.3 ng/mL against infectious virus (Andreano et al., 2020a). Although IC_50_ values measured against pseudovirus correlated well with those measured against infectious virus (*r* = 0.7694, *p* < 0.0001, **Supplemental Figure S3A**), IC_50_ values were typically 5-10 fold less potent against infectious virus as seen previously with patient sera (Seow et al., 2020). Some S1/non-RBD nAbs that had shown weak neutralization (>10 μg/mL) against pseudovirus were only able to neutralize infectious virus at very high concentrations (>50 μg/mL) or undetectable neutralization. In contrast, NTD-specific mAbs P008_056, P008_007 and P003_027 had ~10-fold higher potency against infectious virus (**Supplemental Table S1 and Supplementary Figures 3C and 3D**). In particular, P008_056 neutralized infectious virus with an IC_50_ of 14 ng/mL making this one of the most potent NTD nAbs reported thus far (McCallum et al., 2021). nAbs specific for the RBD generally displayed more potent neutralization compared to those binding non-RBD epitopes (**Figure 4B-C**). Low neutralization plateaus and shallow neutralization curves were observed for some mAbs (Liu et al., 2020) (**Supplementary Figures 3B and 3C**) suggesting incomplete neutralization. These mAbs were typically less potent and had higher IC_50_ values >1000 ng/mL. Seven of the non-S1 binding mAbs bound S2 in ELISA (**Supplemental Table 1**) but none showed neutralizing activity.

**Figure 4:**
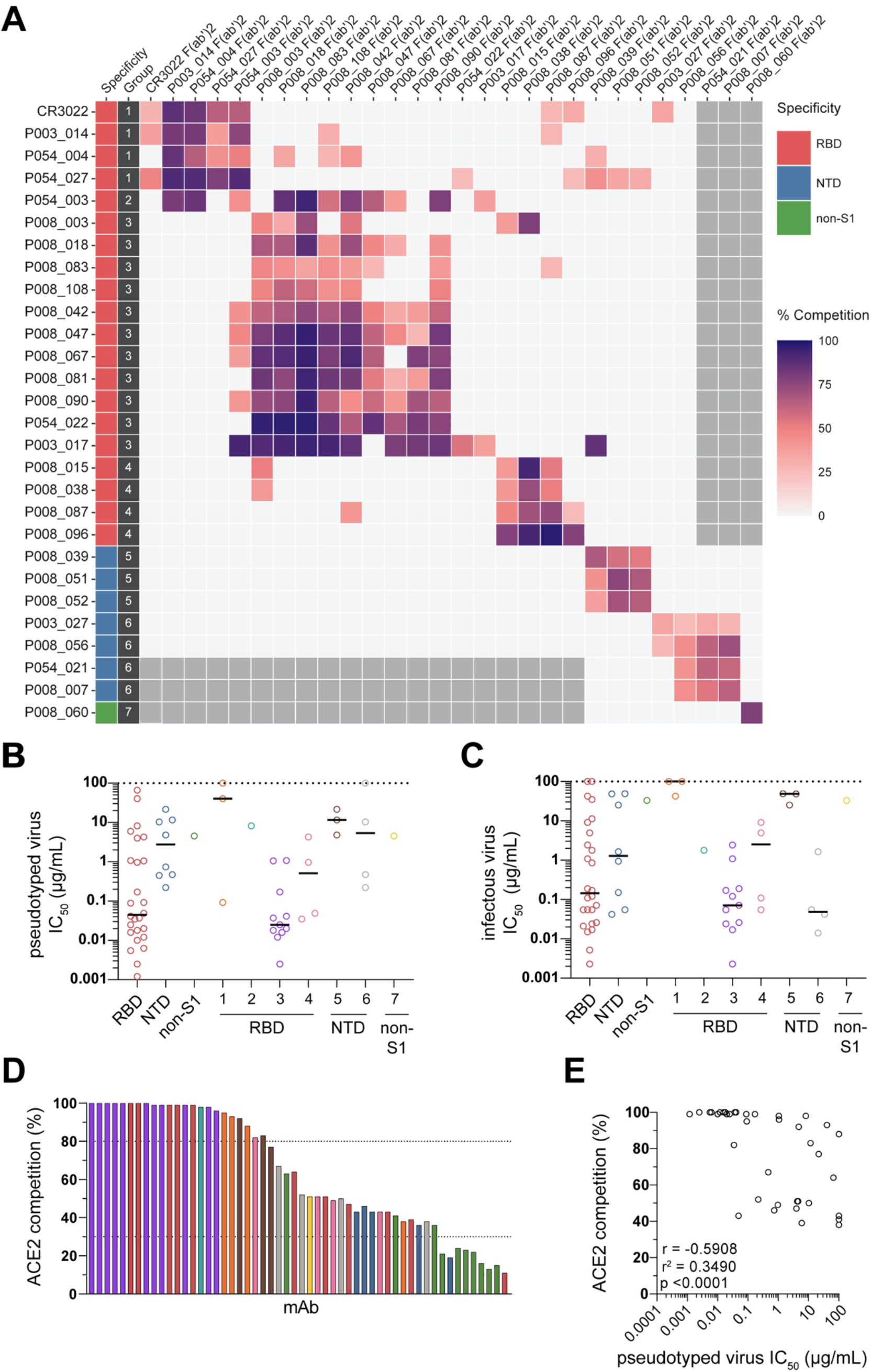
Characterization of SARS-CoV-2 specific neutralizing and non-neutralizing antibodies. **A)** Inhibition of IgG binding to SARS-CoV-2 Spike by F(ab)_2_’ fragments (generated through IdeS digestion of purified IgG) was measured. Serial dilutions of F(ab’)_2_ (starting at 100-molar excess of the IC_80_ of Spike binding) were incubated for 1 hr before washing and addition of competing IgG (added at the IC_80_ of Spike binding). The percentage competition was calculated using the reduction in IgG binding in the presence of F(ab’)_2_ (at 100-molar excess of the IC_80_) as a percentage of the maximum IgG binding in the absence of F(ab’)_2_. Competition groups were determined using Ward2 clustering for initial analysis and clusters were then arranged by hand according to binding epitopes. Competition <25% is white. Grey boxes indicate competition not tested. Neutralization potency (IC_50_) of mAbs targeting either RBD, NTD or non-S1 and/or in competition groups 1-7 against **B)** pseudovirus and **C)** infectious virus. **D)** Ability of neutralizing and non-neutralizing mAbs to inhibit the interaction between cell surface ACE2 and soluble SARS-CoV-2 Spike. mAbs (at 600 nM) were pre-incubated with fluorescently labelled Spike before addition to HeLa-ACE2 cells. The percentage reduction in mean fluorescence intensity is reported. Bars are colour coded based on competition group or Spike subdomain specificity if competition group was not determined. Group 1 (orange), Group 2 (turquoise), Group 3 (purple), Group 4 (pink), Group 5 (brown), Group 6 (grey), Group 7 (yellow), un-mapped RBD (red), un-mapped NTD (blue), S2 (green). **E)** Correlation between IC_50_ against pseudovirus (x-axis) and % ACE2 competition (y-axis). (Spearman correlation, *r*. A linear regression was used to calculate the goodness of fit, *r*^2^).

### Neutralizing antibodies form seven binding competition groups

To gain further insight into epitopes targeted by the isolated mAbs, we performed competition Spike ELISAs between 27 IgG and F(ab)_2_’ fragments (generated through IdeS digestion of purified IgG). Seven distinct competition groups were observed (**Figure 4A**). nAbs binding RBD could be separated into four competition Groups. Group 1 nAbs competed with a previously described SARS-CoV nAb CR3022 which binds an RBD site distal to the ACE2 receptor binding site (Yuan et al., 2020b; Yuan et al., 2020c) (classified Class 4 by Barnes *et al* (Barnes et al., 2020)). Group 1 nAbs displayed limited neutralization potency, particularly against infectious virus (**Supplemental Table 1**). Group 3 nAbs formed the largest and most potent competition group (**Figure 4B&C**). It contained 57.8% (11/19) of RBD nAbs tested in the competition ELISA, and included the most potent mAb P008_108 (IC_50_ 2.3 ng/mL against infectious virus). mAbs in this group were dominated (7/11) by the VH3-53 and VH3-66 germlines (**Supplemental Table 1**) and predominantly clustered together in the phylogenetic tree (**Figure 3A**). A similar VH3-53/VH3-66 gene enrichment has been reported for mAbs that directly bind the ACE2 receptor binding motif (RBM) on RBD (Barnes et al., 2020; Robbiani et al., 2020; Yuan et al., 2020a). Group 2 contained a single nAb with a CDRH3 of 22 amino acids that competed with antibodies in both Groups 1 and 3 suggesting an epitope overlapping these two competition groups. Group 4 contained four RBD reactive mAbs that formed a distinct competition group indicating a further distal RBD neutralizing epitope.

Non-RBD binding nAbs formed three competition groups. Group 5 contained three nAbs which bound Spike, S1 and NTD and had limited neutralization potency (IC_50_ 4.8 – 21.7 μg/mL and 25.3 – 48.8 μg/mL against pseudovirus and infectious virus, respectively). Group 6 contained four nAbs which were all more potent against infectious virus compared to pseudovirus (**Figure 4B&C**) and included P008_056 which neutralized infectious virus with an IC_50_ (14ng/mL) in line with the most potent RBD binding nAbs. Structural analysis revealed that P008_056 binds NTD adjacent to the β-sandwich fold (Rosa et al., 2021). Group 7 contained only one nAb, P008_060, which bound to Spike and not individual S1 or S2 domains suggesting it may target a quaternary epitope spanning multiple domains, similar to 2-43 (Liu et al., 2020).

### mAbs inhibit Spike-ACE2 interaction to differing extents

To explore the potential mechanism by which nAbs prevent infection of target cells, we measured the ability of nAbs to prevent the interaction of Spike with the ACE2 receptor on HeLa cells by flow cytometry (**Figure 4D**). Group 3 nAbs showed >99% inhibition of Spike binding to HeLa-ACE2 cells suggesting that these nAbs directly target the ACE2 binding site on RBD. Overall, nAbs displaying the highest competition with ACE2 binding typically had the highest neutralization potency (**Figure 4E**). Similar to CR3022, Group 1 nAbs showed less complete competition (88.2-95.1%) and Group 4 mAbs show only partial competition (43.1-82.2%) suggesting that these nAbs can sterically inhibit the interaction of Spike with ACE2 without directly binding to the RBM or cause conformational changes to Spike that limit ACE2 binding. NTD-binding and Spike-specific nAbs (Groups 5, 6 and 7) also showed some partial competition (38.4-91.8%). Although these nAbs do not compete with RBD nAbs, it is possible binding to Spike causes conformational changes that prevent subsequent ACE2 binding or lock RBD in the “down” conformation which occludes access to the ACE2 binding site (Liu et al., 2020). As might be expected, S2-reactive mAbs and S1-reactive non-neutralizing mAbs showed negligible competition with ACE2 binding. Overall, these results suggest that some antibodies described here neutralize SARS-CoV-2 through mechanisms beyond direct receptor binding inhibition, such as inhibiting membrane fusion (McCallum et al., 2021) or S1 shedding (Piccoli et al., 2020), which need be investigated further.

### Glycan heterogeneity influences neutralization potency

As mentioned above, some nAbs displayed shallow neutralization curves that plateau before 100% neutralization against pseudovirus and this was more typical for mAbs specific for NTD. Similar unusual neutralization profiles have been observed for some HIV-1 bnAbs, in particular those that accommodate and/or bind *N*-linked glycans on the HIV-1 Env surface, and are thought to arise due to heterogeneity in glycosylation (Doores and Burton, 2010). This phenotype could be rescued for some HIV-1 bnAbs by altering the composition of Env glycans by expressing virus in the presence of glycosidase inhibitors such as kifunensine (that inhibits the ER-mannosidase I enzyme leading to Man9GlcNAc2 glycans) and swainsonine (that inhibits the Golgi-α-mannosidase II enzyme, leading to truncated complex-type glycans in addition to the naturally occurring high-mannose glycans present).

As NTD is heavily glycosylated (Watanabe et al., 2020), we next investigated whether changes in the glycan structures on Spike, through preparation of pseudovirus in the presence of either kifunensine or swainsonine, could affect neutralization activity. RBD mAbs, P008_015, P008_087, P008_090 and P008_108, were not impacted by alterations in Spike glycan processing (**Figure 5**). However, NTD-specific Group 5 mAbs, P008_039, P008_051 and P008_052, and non-S1 Group 7 mAb, P008_060, showed enhanced neutralization against pseudovirus prepared in the presence of swainsonine where glycan structures will be smaller in size. No change in neutralization was observed against pseudovirus prepared with kifunensine although lower infectivity was noted as previously reported (Yang et al., 2020). These data suggest that glycan structures can affect nAb epitope recognition either through modulating the conformation of Spike or altering the accessibility of nAb epitopes.

**Figure 5:**
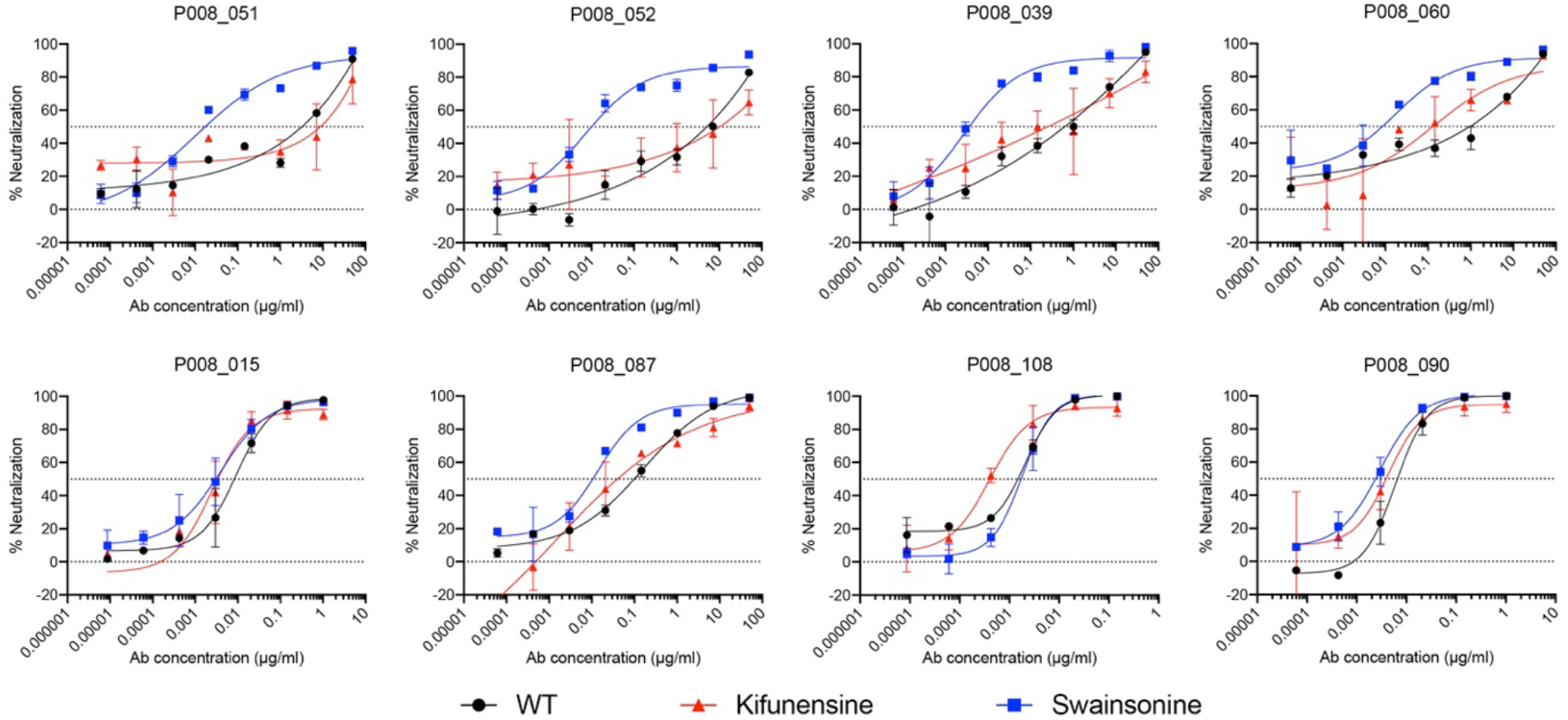
Neutralization of Group 5 nAb enhanced by changes in Spike glycosylation. SARS-CoV-2 pseudovirus was expressed in the presence of glycosidase inhibitors kifunensine or swainsonine. Neutralization potency of RBD and NTD nAbs against Spike modified viruses was measured. Group 5 nAbs (P008_051, P008_052 and P008_039) and Group 7 nAb (P008_060) showed an enhanced neutralization potency and more typical shaped neutralization curve compared to Spike with wild-type glycans. In contrast, RBD targeting nAbs (P008_015, P008_087, P008_108 and P008_090) had unchanged neutralization.

### Cross-reactivity of nAbs with SARS-CoV

SARS-CoV Spike shares 73% sequence homology with Spike of SARS-CoV-2 and 73% with RBD (Walls et al., 2020). To determine whether isolated mAbs targeted epitopes shared between SARS-CoV-2 and SARS-CoV, cross-neutralization against SARS-CoV was measured using the HIV-1 pseudovirus system expressing full-length SARS-CoV Spike. Although neutralization was detected for several mAbs, this was generally at a much-reduced potency (3- to 65-fold) compared to SARS-CoV-2 neutralization (**Supplemental Table 1**). nAb CR3022, isolated following SARS-CoV infection, has previously been reported to bind a conserved epitope between SARS-CoV and SARS-CoV-2 (Yuan et al., 2020c). However, cross-neutralization was observed for only one nAb in the CR3022 competition group (Group 1) and single nAbs from Groups 4 (RBD), 5 (NTD) and 7 (Spike only) suggesting nAbs within each competition group bind the same footprint but differ in their molecular contacts. Interestingly, the sole mAb in Group 7, which reacts only with Spike trimer and not individual subunits, showed a 7-fold more potent neutralization against SARS-CoV compared to SARS-CoV-2. Binding of nAbs to SARS-CoV Spike expressed on the surface of HEK 293T cells was detected for mAbs showing SARS-CoV neutralizing activity but not by SARS-CoV non-neutralizing antibodies in the same competition groups (**Supplementary Figure 3E**). However, S2-binding non-neutralizing antibodies, although unable to neutralize SARS-CoV (**Supplemental Table 1**), bound to cell surface expressed SARS-CoV Spike (**Supplementary Figure 3F**) indicating the presence of a conserved, non-neutralizing S2 epitope. Whether the S2 mAbs can bind SARS-CoV-2 infected cells and recruit ADCC *in vivo* is not known (Li et al., 2020b). Overall, conserved neutralizing epitopes shared between SARS-CoV-2 and SARS-CoV are present on both RBD and NTD.

### Sensitivity to nAbs to newly emerging Spike variants

The SARS-CoV-2 D614G Spike variant supplanted the ancestral virus in most areas worldwide early in the pandemic and although the mutation has been reported to be more infectious through stabilization of the RBD in the ‘up’ conformation, it has not been associated with neutralization escape (Li et al., 2020a; Weissman et al., 2020; Yurkovetskiy et al., 2020). More recently the B.1.1.7 variant of concern first reported in the UK, which contains an additional eight Spike mutations in NTD, RBD and S2 (ΔH69/V70, ΔY144, N501Y, A570D, P681H, T716I, S982A, D1118H) (Rambaut et al., 2020), has been associated with more efficient transmission within the UK and is now the dominant variant in London and the South East of England (Rambaut et al., 2020). It is not known whether these mutations have arisen stochastically, have been selected purely on the basis of increased transmission, or whether the emergence of B.1.1.7 was in part driven by the pressure of neutralizing antibodies in longer term infections in immunocompromised patients undergoing passive immunotherapy (Kemp et al., 2020; Mccarthy et al., 2020). Nor is it clear if it will lead to escape from the neutralizing antibodies generated in response to SARS-CoV-2 infection in wave 1 and/or generated through vaccination. Initial reports have suggested that the B.1.1.7 variant is sensitive to polyclonal sera from individuals infected with early circulating SARS-CoV-2 variants (Rees-Spear et al., 2021).

We measured neutralization potency of nAbs from the seven competition groups, as well as patient plasma from P008 and P054, against HIV-1 viral particles pseudotyped with SARS-CoV-2 Spike bearing mutations i) D614G, ii) N501Y (part of the ACE2 receptor binding site and associated with increased transmission), iii) D614G+ΔH69/V70 (H69 and V70 deletion in NTD loop), and iv) the B.1.1.7 variant containing all eight Spike mutations. Similar to previous data (Yurkovetskiy et al., 2020), the D614G mutation alone only showed a modest effect on neutralization by the majority of RBD-specific nAbs or plasma from the P008 and P054 donors. However, NTD-specific nAbs within competition group 5 showed a decrease (22-140 fold) in neutralization potency against the D614G variant (**Figure 6A** and **Supplemental Figure 4A**) and two group 4 RBD-specific nAbs, P008_087 and P008_038, showed a 4- and 15-fold decrease in potency respectively and Group 1 RBD-specific nAb, P054_027, showed a 16-fold reduction. N501 is part of the ACE2 receptor binding motif. Substitution to tyrosine has been shown to increase affinity for ACE2 (Starr et al., 2021) and the N501Y mutation was observed in a mouse adapted SARS-CoV-2 variant (Gu et al., 2020). Despite the location of the N501Y mutation in RBD, the vast majority of RBD specific nAbs were not affected by this Spike mutation (**Figure 6A** and **Supplemental Figure 4A**). By contrast, group 3 nAbs P003_017 and P008_003 showed a 15- and 6-fold decrease respectively, and group 1 nAb P054_004 showed a 5-fold decrease in neutralization potency against the N501Y variant. No change in neutralization was observed for NTD-specific group 5 and 6 mAbs against the N501Y mutant. The ΔH69/V70 is situated within the N1 loop of NTD and, in addition to being present in the B.1.1.7 variant, has been associated with viral evolution in an immunocompromised SARS-CoV-2 infected individual undergoing convalescent plasma therapy (Kemp et al., 2020). Deletion H69/V70 in combination with the D614G mutation had a very limited effect on neutralization of RBD-specific or NTD-specific antibodies with only P054_004 and P003_027 showing very small reductions (7-8 fold) compared to the D614G variant (**Figure 6B** and **Supplemental Figure 4B**).

**Figure 6:**
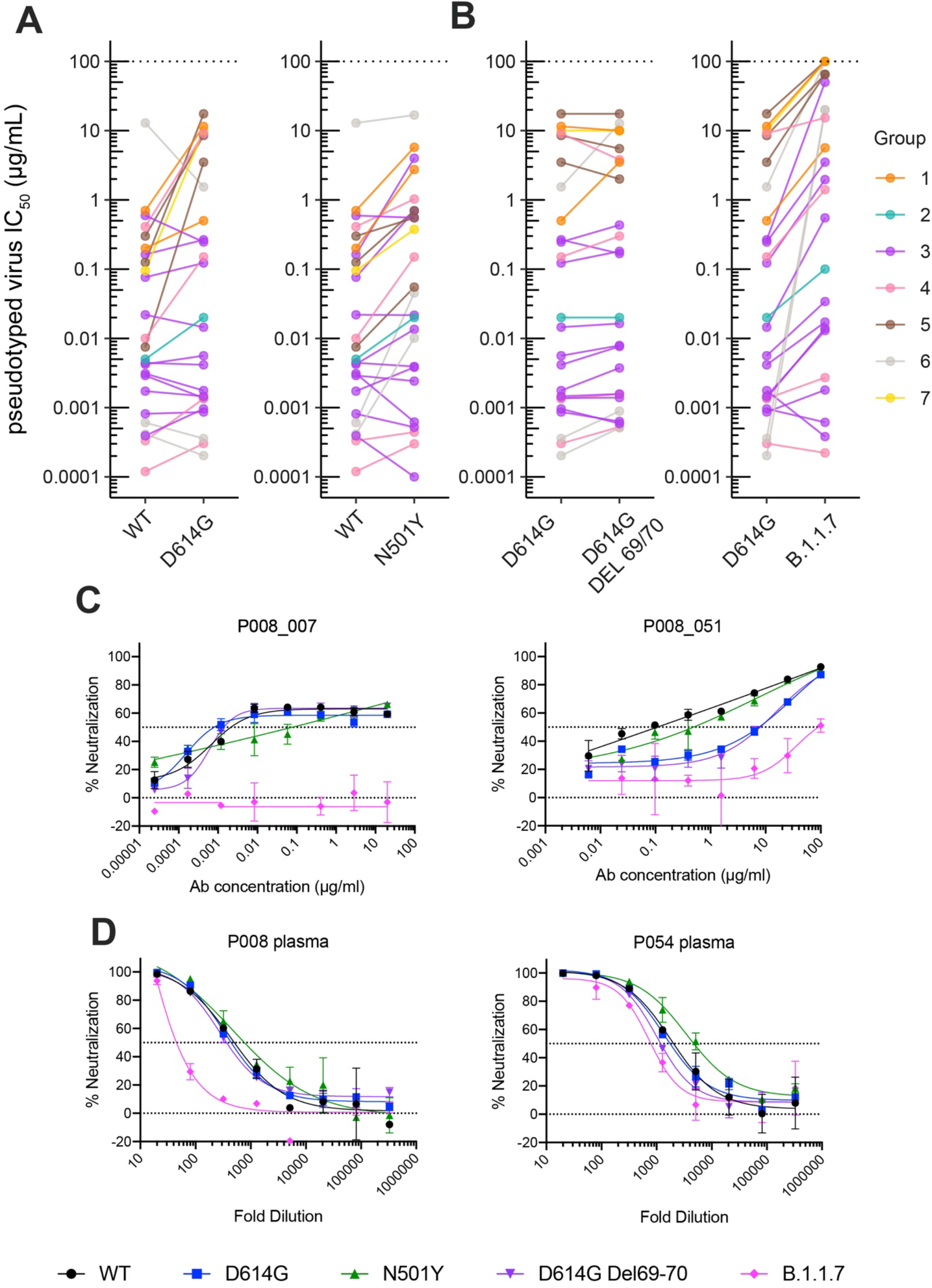
Susceptibility of B.1.1.7 and variants to neutralization by neutralizing antibodies. Neutralization of mAbs and plasma were tested against pseudoviruses expressing variant Spikes including D614G, N501Y and D614G ΔH69/V70 mutations and the B.1.1.7 variant. **A)** Change in neutralization IC_50_ (μg/mL) for D614G and N501Y mutation compared to wild-type Spike. nAbs are coloured by competition group according to the key. **B)** Change in neutralization IC_50_ (μg/mL) for D614G ΔH69/V70 mutation and B.1.1.7 variant compared D614G Spike. **C)** Example neutralization curves for a group 5 and group 6 NTD-specific nAb against Spike variants. **D)** Neutralization activity in plasma from P008 and P054 at the sorting time point against Spike variants.

NTD nAbs in both group 5 and 6 showed a dramatic reduction in neutralization against the B.1.1.7 variant (**Figure 6B-D** and **Supplemental Figure 4B**). Group 6 NTD-specific nAbs (including P008_056, P008_007, P003_027 and P054_021) showed no detectable neutralizing activity up to 100 μg/mL and group 5 NTD-specific nAbs showed a >200-fold reduction in potency. The most potent mAbs in RBD groups 3 and 4 showed no reduction in neutralization activity against the B.1.1.7 variant. In contrast, P008_081, P054_022 and P008_047 (Group 3) and P008_096 and P008_087 (Group 4) showed a 4 – 38-fold reduction against B.1.1.7 whilst maintaining IC_50_ values in the 0.013-15 μg/mL range. P003_017 was the RBD nAb showing the greatest reduction in neutralization with only very weak neutralization detected at 50 μg/mL. Despite the reduction in neutralization potency or loss of neutralization activity for specific mAbs, neutralization potency of sera from P008 and P054 at days 61 and 48 POS showed a minimal 2-fold and 8-fold reduction in potency against the B.1.1.7 variant (**Figure 6D**), respectively, suggesting the polyclonal nature of the antibody response overcomes the Spike mutations in the B.1.1.7 variant.

## Discussion

Here we isolated SARS-CoV-2 Spike reactive neutralizing and non-neutralizing antibodies from three convalescent donors who experienced a range of COVID-19 illness severities. As observed previously (Barnes et al., 2020; Brouwer et al., 2020; Jiang et al., 2020; Liu et al., 2020; Pinto et al., 2020; Robbiani et al., 2020; Rogers et al., 2020; Tortorici et al., 2020; Wu et al., 2020), the antibody response to SARS-CoV-2 is very diverse, is not restricted to specific germlines and does not require extensive somatic hypermutation for neutralization as seen for HIV-1 nAbs (Doores et al., 2015; Scheid et al., 2009). Antibodies against RBD, NTD and non-S1 epitopes were isolated from all three donors. The most potent neutralizing antibody, P008_108 (IC_50_ 2.3 ng/mL against infectious virus), was isolated from an asymptomatic donor with very low serum neutralizing activity. Isolation of similar potent mAbs from donors with low serum neutralizing activity has also been reported by Robbiani *et al* (Robbiani et al., 2020) and suggests that the lower neutralization potency of sera from individuals experiencing mild or no COVID-19 symptoms is due to a low quantity of plasma neutralizing antibody rather than sub-optimal potency of individual antibodies. It also suggests that memory responses are not proportional to the antibodies arising in serum after the immediate plasmablast burst. Upon re-exposure, all individuals would be expected to produce highly potent neutralizing antibodies.

The use of Spike for antigen-specific B cell sorting allowed us to isolate mAbs targeting epitopes beyond the RBD and to study their relevance in viral evolution and antigenic changes in Spike. Roughly one third of isolated mAbs were specific for NTD but only 28.5% of these showed neutralizing activity. Neutralizing NTD mAbs formed two distinct competition groups (**Figure 4A**). The most potent competition group (Group 6) contained nAbs that neutralized infectious virus more potently than pseudovirus. Assuming the structural conformations and dynamics of the Spike on pseudovirus and infectious virus are identical, the discordant neutralization differences may relate to differences in Spike density, Spike glycan heterogeneity or levels of the ACE2 receptor on the target cell lines used in the two assays. nAb P008_056 neutralized infectious virus with an IC_50_ of 14 ng/mL making this one of the most potent NTD nAbs reported (Rogers et al., 2020) and in line with the most potent RBD nAb isolated here. Structural analysis of the interaction of P008_056 Fab with Spike revealed that the antibody binds the viral glycoprotein at the distal face of the NTD, including dramatic conformation changes in this domain (Rosa et al., 2021). The less potent NTD competition group (Group 5) showed atypical neutralization curves. For these antibodies, neutralization activity could be enhanced by reducing the size and/or composition of *N*-glycans on Spike through preparation of pseudoviral particles in the presence of swainsonine (**Figure 5**). As NTD is heavily glycosylated, a reduction in the glycan size will likely increase the accessibility of protein epitopes on NTD (Watanabe et al., 2020) and thus enhance binding efficiency and neutralization potency.

RBD-specific nAbs isolated here targeted epitopes similar to those reported previously (Barnes et al., 2020; Brouwer et al., 2020; Piccoli et al., 2020; Seydoux et al., 2020; Yuan et al., 2020b; Zost et al., 2020) and included nAbs that directly block ACE2 binding through binding RBM and nAbs targeting epitopes distal to the RBM. We also isolated a high proportion of Spike reactive mAbs which showed no neutralizing activity demonstrating the presence of immunodominant, non-neutralizing epitopes on RBD, NTD and S2. 31.8% (34/107) of mAbs did not bind suggesting S2 or quaternary epitopes, but only three showed neutralizing activity. Although S2 reactive mAbs showed no neutralizing activity, they were able to cross-react with SARS-CoV Spike expressed on the surface of cells. As the non-neutralizing mAbs are able to bind cell-surface expressed Spike, it will be important to investigate whether they can facilitate Fc effector functions such as ADCC and play a role in virus clearance (Bournazos et al., 2020; Li et al., 2020b).

New SARS-CoV-2 variants are rapidly emerging across the globe and it is important to determine whether antibodies generated during wave 1 infections or following vaccination will provide protection against these new variants of concern (Rambaut et al., 2020). Despite the dominant neutralizing antibody response being against RBD, we show that the B.1.1.7 variant is still potently neutralized by some RBD-specific nAbs but is highly resistant to NTD-specific nAbs. As the ΔH69/V70 deletion alone did not lead to neutralization resistance, it is likely the ΔY144 deletion facilitates neutralization escape. Indeed, structural analysis of P008_056 in complex with SARS-CoV-2 Spike shows that Y144 sits within a loop that must undergo conformational rearrangement to allow access to the P008_056 epitope on NTD (Rosa et al., 2021). It is not possible to conclude from our data whether nAbs against NTD are selecting for Spike variants encoding NTD deletions, such as B.1.1.7, or whether NTD mutations alter Spike functionality to favour increased transmissibility. More specifically, deletions in NTD have been associated with neutralization escape from mAbs (Andreano et al., 2020b; Kemp et al., 2020; Mccarthy et al., 2020; Wimber et al., 2021). The ΔY144 deletion, which has a prevalence of 1.7% in 37 countries (McCallum et al., 2021), has been shown to abrogate binding to other NTD mAbs including S2M28, S2X28, S2X333 and 4A8 (Chi et al., 2020; McCallum et al., 2021). NTD deletions, including ΔH69/V70 (Kemp et al., 2020) and Δ141-144 (Avanzato et al., 2020; Choi et al., 2020; Mccarthy et al., 2020), have been observed in immunocompromised individuals who remain infected for extended periods. And lastly, deletion of NTD residues 242-244 from the B.1.351 variant (501Y.V2 prevalent in South Africa) and has been shown to reduce binding by NTD specific mAbs 4A8 (Wimber et al., 2021) and 4-8 (P. et al., 2021). Thus, more research is needed to establish the driver for the observed accumulation of genetic changes in circulating SARS-CoV-2 strains.

Despite the loss in neutralization of NTD-specific nAbs against B.1.1.7, neutralization by RBD-specific nAbs either remained unchanged or, when a reduction was observed, neutralization in the 0.001 – 5 μg/mL range was still measured for the majority of antibodies. The reduction in RBD-specific mAbs neutralization against B.1.1.7 was of lower magnitude than that reported for mAbs against the B.1.351 variant (Wimber et al., 2021). The B.1.351 variant also encodes RBD mutations K417N and E484K which have been associated with viral escape from RBD-targeting antibodies (Starr et al., 2021; Weisblum et al., 2020). As RBD is the predominant target for neutralizing antibodies following infection (Greaney et al., 2020; Piccoli et al., 2020), this would suggest RBD-specific nAbs had a limited contribution to any immune escape contributing to the selection of the B.1.1.7 variant. Importantly, although there was a 2- and 8-fold decrease in P054 and P008 plasma neutralization, respectively, against the B.1.1.7 variant, neutralization could still be detected. This suggests that although complete loss of neutralization is observed for specific mAbs, further mutations would be needed for complete neutralization escape from the polyclonal antibody response generated from SARS-CoV-2 infection in these individuals.

In conclusion, we identified potent neutralizing antibodies targeting both RBD and NTD neutralizing epitopes. We show that the B.1.1.7 variant is resistant to neutralization by the NTD nAbs demonstrating the importance of considering both dominant and sub-dominant neutralizing epitopes on Spike when studying viral evolution and antigenic drift.

## Methods

### Ethics statement

This study used samples collected as part of the COVID-IP study.(Laing et al., 2020a) The study protocol for patient recruitment and sampling, out of the intensive care setting, was approved by the committee of the Infectious Diseases Biobank of King’s College London with reference number COV-250320. The protocol for healthy volunteer recruitment and sampling was similarly approved by the same committee as an amendment to an existing approval for healthy volunteer recruitment with reference number MJ1-031218b. Both approvals were granted under the terms of the Infectious Disease Biobank’s ethics permission (reference 19/SC/0232) granted by the South Central Hampshire B Research Ethics Committee in 2019. Patient recruitment from the ICU was undertaken through the ethics for the IMMERSE study approved by the South Central Berkshire Ethics Committee with reference number 19/SC/0187. Patient and control samples and data were anonymized at the point of sample collection by research nursing staff or clinicians involved in the COVID-IP project. We complied with all relevant ethical regulations.

### Protein expression and purification

Recombinant Spike and RBD for ELISA were expressed and purified as previously described (Seow et al., 2020). Recombinant S1 (residues 1-530) and NTD (residues 1-310) expression and purification was described in Rosa et al (Rosa et al., 2021). S2 protein was obtained from SinoBiological (Cat number: 40590-V08B).

For antigen-specific B cell sorting, Spike glycoprotein consisted of the pre-fusion S ectodomain (residues 1–1138) with a GGGG substitution at the furin cleavage site (amino acids 682–685), proline substitutions at amino acid positions 986 and 987, and an N-terminal T4 trimerization domain. Spike was cloned into a pHLsec vector containing Avi and 6xHis tags (Aricescu et al., 2006). Biotinylated Spike was expressed in 1L of HEK293F cells (Invitrogen) at a density of 1.5 × 10^6^ cells ml^-1^. To achieve in-vivo biotinylation, 480μg of each plasmid was co-transfected with 120μg of BirA and 12mg PEI-Max (Polysciences) in the presence of 200μM biotin (final concentration). The supernatant was harvested after 7 days and purified using immobilized metal affinity chromatography and size-exclusion chromatography. Complete biotinylation was confirmed via depletion of proteins using avidin beads.

### ELISA (S, RBD, NTD, S2 or S1)

96-well plates (Corning, 3690) were coated with S, S1, NTD, S2 or RBD at 3μg/mL overnight at 4°C. The plates were washed (5 times with PBS/0.05% Tween-20, PBS-T), blocked with blocking buffer (5% skimmed milk in PBS-T) for 1 h at room temperature. Serial dilutions of serum, plasma, mAb or supernatant in blocking buffer were added and incubated for 2 hr at room temperature. Plates were washed (5 times with PBS-T) and secondary antibody was added and incubated for 1 hr at room temperature. IgM was detected using Goat-anti-human-IgM-HRP (horseradish peroxidase) (1:1,000) (Sigma: A6907) and IgG was detected using Goat-anti-human-Fc-AP (alkaline phosphatase) (1:1,000) (Jackson: 109-055-098). Plates were washed (5 times with PBS-T) and developed with either AP substrate (Sigma) and read at 405 nm (AP) or 1-step TMB (3,3’,5,5’-Tetramethylbenzidine) substrate (Thermo Scientific) and quenched with 0.5 M H_2_S0_4_ before reading at 450 nm (HRP).

### Fab/Fc ELISA

96-well plates (Corning, 3690) were coated with goat anti-human Fc IgG antibody at 3 μg/mL overnight at 4°C. The above protocol was followed. The presence of IgG in supernatants was detected using Goat-anti-human-Fc-AP (alkaline phosphatase) (1:1,000) (Jackson: 109-055-098).

### IgG digestion to generate F(ab’)_2_

IgG were incubated with IdeS (4 μg of IdeS per 1 mg of IgG) in PBS for 1 hour at 37 °C. The Fc and IdeS A were removed using a mix of Protein A Sepharose^®^ Fast Flow (250 μL per 1 mg digested mAb; GE Healthcare Life Sciences) and Ni Sepharose™ 6 Fast Flow (50 μL per 1 mg digested mAb; GE Healthcare Life Sciences) which were washed twice with PBS before adding to the reaction mixture. After exactly 10 minutes the beads were removed from the F(ab’)_2_-dilution by filtration in Spin-X tube filters (Costar^®^) and the filtrate was concentrated in Amicon^®^ Ultra Filters (10k, Millipore). Purified F(ab’)_2_ fragments were analysed by SDS-PAGE.

### F(ab’)_2_ and IgG competition ELISA

96-well half area high bind microplates (Corning^®^) were coated with S-protein at 3μg/mL in PBS overnight at 4 °C. Plates were washed (5 times with PBS/0.05% Tween-20, PBS-T) and blocked with 5% milk in PBS/T for 2 hr at room temperature. Serial dilutions (5-fold) of F(ab’)_2_, starting at 100-molar excess of the IC_80_ of S binding, were added to the plates and incubated for 1 hr at room temperature. Plates were washed (5 times with PBS-T) before competing IgG was added at their IC_80_ of S binding and incubated at room temperature for 1 hr. Plates were washed (5 times with PBS-T) and Goat-anti-human-Fc-AP (alkaline phosphatase) (1:1,000) (Jackson: 109-055-098) was added and incubated for 45 minutes at room temperature. The plates were washed (5 times with PBS-T) and AP substrate (Sigma) was added. Optical density was measured at 405 nm in 5-minute intervals. The percentage competition was calculated as the reduction in IgG binding in the presence of F(ab’)_2_ (at 100-molar excess of the IC_80_) as a percentage of the maximum IgG binding in the absence of F(ab’)_2_. Competition groups were determined using Ward2 clustering (R, Complex Heatmap package (Gu et al., 2016)) for initial analysis and Groups were then arranged by hand according to binding epitopes.

### SARS-CoV-2 (wild-type and mutants) and SARS-CoV pseudotyped virus preparation

Pseudotyped HIV virus incorporating the SARS-Cov-2 wild-type or mutants (D614G, N501Y, D614G+Del69/70 and B.1.1.7) or SARS-CoV spike protein was produced in a 10 cm dish seeded the day prior with 5×10^6^ HEK293T/17 cells in 10 ml of complete Dulbecco’s Modified Eagle’s Medium (DMEM-C, 10% foetal bovine serum (FBS) and 1% Pen/Strep (100 IU/ml penicillin and 100 mg/ml streptomycin)). Cells were transfected using 90 mg of PEI-Max (1 mg/mL, Polysciences) with: 15 μg of HIV-luciferase plasmid, 10 μg of HIV 8.91 gag/pol plasmid and 5 μg of SARS-CoV-2 spike protein plasmid (Grehan et al., 2015; Thompson, 2020). Pseudotyped virus was harvested after 72 hours, filtered through a 0.45mm filter and stored at −80°C until required.

### Viral entry inhibition assay with SARS-CoV-2 (wild-type and mutants) and SARS-CoV pseudotyped virus

Neutralization assays were conducted as previously described (Carter et al., 2020). Serial dilutions of serum samples (heat inactivated at 56°C for 30mins) or mAbs were prepared with DMEM-C media and incubated with pseudotyped virus for 1-hour at 37°C in 96-well plates. Next, HeLa cells stably expressing the ACE2 receptor (provided by Dr James Voss, Scripps Research, La Jolla, CA) were added (12,500 cells/50μL per well) and the plates were left for 72 hours. Infection level was assessed in lysed cells with the Bright-Glo luciferase kit (Promega), using a Victor™ X3 multilabel reader (Perkin Elmer). Measurements were performed in duplicate and duplicates used to calculate the ID_50_.

### Infectious virus strain and propagation

Vero-E6 cells (Cercopithecus aethiops derived epithelial kidney cells, provided by Prof Wendy Barclay, Imperial College London) cells were grown in Dulbecco’s modified Eagle’s medium (DMEM, Gibco) supplemented with GlutaMAX, 10% fetal bovine serum (FBS), 20 μg/mL gentamicin, and incubated at 37°C with 5% CO_2_. SARS-CoV-2 Strain England 2 (England 02/2020/407073) was obtained from Public Health England. The virus was propagated by infecting 60-70% confluent Vero E6 cells in T75 flasks, at an MOI of 0.005 in 3 ml of DMEM supplemented with GlutaMAX and 10% FBS. Cells were incubated for 1 hr at 37°C before adding 15 ml of the same medium. Supernatant was harvested 72h post-infection following visible cytopathic effect (CPE), and filtered through a 0.22 μm filter to eliminate debris, aliquoted and stored at −80C. The infectious virus titre was determined by plaque assay using Vero E6 cells.

### Infectious virus neutralization assay

Vero-E6 cells were seeded at a concentration of 20,000 cells/100uL per well in 96-well plates and allowed to adhere overnight. Serial dilutions of mAbs were prepared with DMEM media (2% FBS and 1% Pen/Strep) and incubated with authentic SARS-CoV-2 for 1 hour at 37°C. The media was removed from the pre-plated Vero-E6 cells and the serum-virus mixtures were added to the Vero E6 cells and incubated at 37°C for 24 h. The virus/serum mixture was aspirated, and each well was fixed with 150μL of 4% formalin at room temperature for 30 min and then topped up to 300μL using PBS. The cells were washed once with PBS and permeabilised with 0.1% Triton-X in PBS at room temperature for 15 min. The cells were washed twice with PBS and blocked using 3% milk in PBS at room temperature for 15 min. The blocking solution was removed and an N-specific mAb (murinized-CR3009) was added at 2μg/mL (diluted using 1% milk in PBS) at room temperature for 45 min. The cells were washed twice with PBS and horse-anti-mouse-IgG-conjugated to HRP was added (1:2000 in 1% milk in PBS, Cell Signaling Technology, S7076) at room temperature for 45 min. The cells were washed twice with PBS, developed using TMB substrate for 30 min and quenched using 2M H_2_SO_4_ prior to reading at 450 nm. Measurements were performed in duplicate.

### Antigen-specific B cell sorting

Fluorescence-activated cell sorting of cryopreserved PBMCs was performed on a BD FACS Melody. Sorting baits (SARS-CoV-2 Spike) was pre-complexes with the streptavidin flurophore at a 1:4 molar ratio prior to addition to cells. PBMCs were stained with live/dead (fixable Aqua Dead, Thermofisher), anti-CD3-APC/Cy7 (Biolegend), anti-CD8-APC-Cy7 (Biolegend), anti-CD14-BV510 (Biolegend), anti-CD19-PerCP-Cy5.5 (Biolegend), anti-IgM-PE (Biolegend), anti-IgD-Pacific Blue (Biolegend) and anti-IgG-PeCy7 (BD) and Spike-Alexa488 (Thermofisher Scientific, S32354) and Spike-APC (Thermofisher Scientific, S32362). Live CD3/CD8^-^CD14^-^CD19^+^IgM^-^IgD^-^IgG^+^Spike^+^Spike^+^ cells were sorted into individual wells containing RNase OUT (Invitrogen), First Strand SuperScript III buffer, DTT and H_2_O (Invitrogen) and RNA was converted into cDNA (SuperScript III Reverse Transcriptase, Invitrogen) using random hexamers following the manufacturer’s protocol.

### Full-length antibody cloning and expression

The human Ab variable regions of heavy and kappa/lambda chains were PCR amplified using previously described primers and PCR conditions (Scheid et al., 2009; Tiller et al., 2008; von Boehmer et al., 2016). PCR products were purified and cloned into human-IgG (Heavy, Kappa or Lambda) expression plasmids(von Boehmer et al., 2016) using the Gibson Assembly Master Mix (NEB) following the manufacturer’s protocol. Gibson assembly products were directly transfected into HEK-293T cells and transformed under ampicillin selection. Ab supernatants were harvested 3 days after transfection and IgG expression and Spike-reactivity determined using ELISA and concentrated 30-times for use in neutralization assays. Ab variable regions of heavy-light chain pairs that generated Spike reactive IgG were sequenced by Sanger sequencing.

### IgG expression and purification

Ab heavy and light plasmids were co-transfected at a 1:1 ratio into HEK-293F cells (Thermofisher) using PEI Max 40K (1mg/mL, linear polyethylenimine hydrochloride, Polysciences, Inc.) at a 3:1 ratio (PEI max:DNA). Ab supernatants were harvested five days following transfection, filtered and purified using protein G affinity chromatography following the manufacturer’s protocol (GE Healthcare).

### Monoclonal antibody binding to Spike using flow cytometry

HEK293T cells were plated in a 6-well plate (2×10^6^ cells/well). Cells were transfected with 1 μg of plasmid encoding fulllength SARS-CoV or SARS-CoV-2 full-length Spike and incubated for 48h at 37°C. Post incubation cells were resuspended in PBS and plated in 96-well plates (1×10^5^ cells/well). Monoclonal antibodies were diluted in FACS buffer (1x PBS, 2% FBS, 1 mM EDTA) to 25 μg/mL and incubated with cells on ice for 1h. The plates were washed twice in FACS buffer and stained with 50 μl/well of 1:200 dilution of PE-conjugated mouse anti-human IgG Fc antibody (BioLegend) on ice in dark for 1h. After another two washes, stained cells were analyzed using flow cytometry, and the binding data were generated by calculating the percent (%) PE-positive cells using FlowJo 10 software.

### ACE2 competition measured by flow cytometry

To prepare the fluorescent probe, 3.5 times molar excess of Streptavadin-PE (Thermofisher Scientific, S21388) was added to biotinylated SARS-CoV-2 Spike on ice. Additions were staggered over 5 steps with 30 min incubation times between each addition. Purified mAbs were mixed with PE conjugated SARS-CoV-2 S in a molar ratio of 4:1 in FACS buffer (2% FBS in PBS) on ice for 1 h. HeLa-ACE2 and HeLa cells were washed once with PBS and detached using PBS containing 5mM EDTA. Detached cells were washed and resuspended in FACS buffer. 0.5 million HeLa-ACE2 cells were added to each mAb-Spike complex and incubated on ice for 30 m. The cells were washed with PBS and resuspended in 200 μL FACS buffer with 1 μL of LIVE/DEAD Fixable Aqua Dead Cell Stain Kit (Invitrogen). HeLa and HeLa-ACE2 cells alone and with SARS-CoV-2 Spike only were used as background and positive controls, respectively. The geometric mean fluorescence for PE was measured from the gate of singlet and live cells. The ACE2 binding inhibition percentage was calculated with this equation: (Rogers et al., 2020)

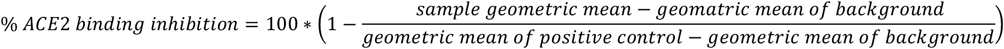

### Sequence analysis/tree generation

The heavy and light chain sequences of SARS-CoV-2 specific mAbs were examined using IMGT/V-QUEST(http://www.imgt.org/IMGT_vquest/vquest) to identify the germline usages, level of SHM and CDR region lengths. To remove variation introduced through cloning using mixture of forward primers, 5 amino acids or 15 nucleotides were trimmed from the start and end of the translated variable genes. The sequences were aligned with Clustal W and clustered via PhyML to produce maximum likelihood phylogenetic trees which were visualised and annotated using FigTree. The Sankey plot was made in Tableau.

## Funding

This work was funded by; King’s Together Rapid COVID-19 Call awards to MHM, KJD and SJDN, MRC Discovery Award MC/PC/15068 to SJDN, KJD and MHM, Fondation Dormeur, Vaduz for funding equipment to KJD, Huo Family Foundation Award to MHM, KJD, MSH and SJDN, MRC Programme Grant (MR/S023747/1 to MHM), Wellcome Trust Multi-User Equipment Grant 208354/Z/17/Z to MHM, SJDN, KD and ACH, Wellcome Trust Investigator Award 106223/Z/14/Z to MHM, NIAID Award U54 AI150472 to MHM, The UK CIC (Covid-Immunology-Consortium) to ACH, the Rosetrees Trust to ACH, and UCL Coronavirus Response Fund to LEM (made possible through generous donations from UCL’s supporters, alumni, and friends). MSH is funded by the National Institute for Health Research Clinician Scientist Award (CS-2016-16-011). The views expressed in this publication are those of the author(s) and not necessarily those of the NHS, the National Institute for Health Research or the Department of Health and Social Care. CG and HW was supported by the MRC-KCL Doctoral Training Partnership in Biomedical Sciences (MR/N013700/1). SA was supported by an MRC-KCL Doctoral Training Partnership in Biomedical Sciences industrial Collaborative Award in Science & Engineering (iCASE) in partnership with Orchard Therapeutics (MR/R015643/1). Work in PC laboratory is funded by the Francis Crick Institute (FC001061), which receives its core funding from Cancer Research UK, the UK Medical Research Council, and the Wellcome Trust. T.A.B. is supported by the Medical Research Council (MR/S007555/1). The Wellcome Centre for Human Genetics is supported by Wellcome Centre grant 203141/Z/16/Z. LEM was supported by a Medical Research Council Career Development Award (MR/R008698/1). This work was supported by the Department of Health via a National Institute for Health Research comprehensive Biomedical Research Centre award to Guy’s and St. Thomas’ NHS Foundation Trust in partnership with King’s College London and King’s College Hospital NHS Foundation Trust.

Thank you to Florian Krammer for provision of the RBD expression plasmid, Philip Brouwer, Marit Van Gils and Rogier Sanders (University of Amsterdam) for the Spike protein construct, and James Voss and Deli Huang for providing the Hela-ACE2 cells.

