## Supplemental information for "Impact of the B.1.1.7 variant on neutralizing monoclonal antibodies recognizing diverse epitopes on SARS-CoV-2 Spike"

**Supplementary figures 1-4**

**Supplementary table 1**

**Supplementary Figure S1: Example sorting strategy to isolate SARS-CoV-2 Spike specific $IgG^+$ B cells.** Figure showing example sorting for donor P008. Live $CD3/CD8^-CD14^-$ $CD19^+IgM^-IgD^-IgG^+Spike^+Spike^+$ cells were sorted into individual wells. The heavy and light chains were reverse transcribed and amplified using nested PCR with gene specific primers.

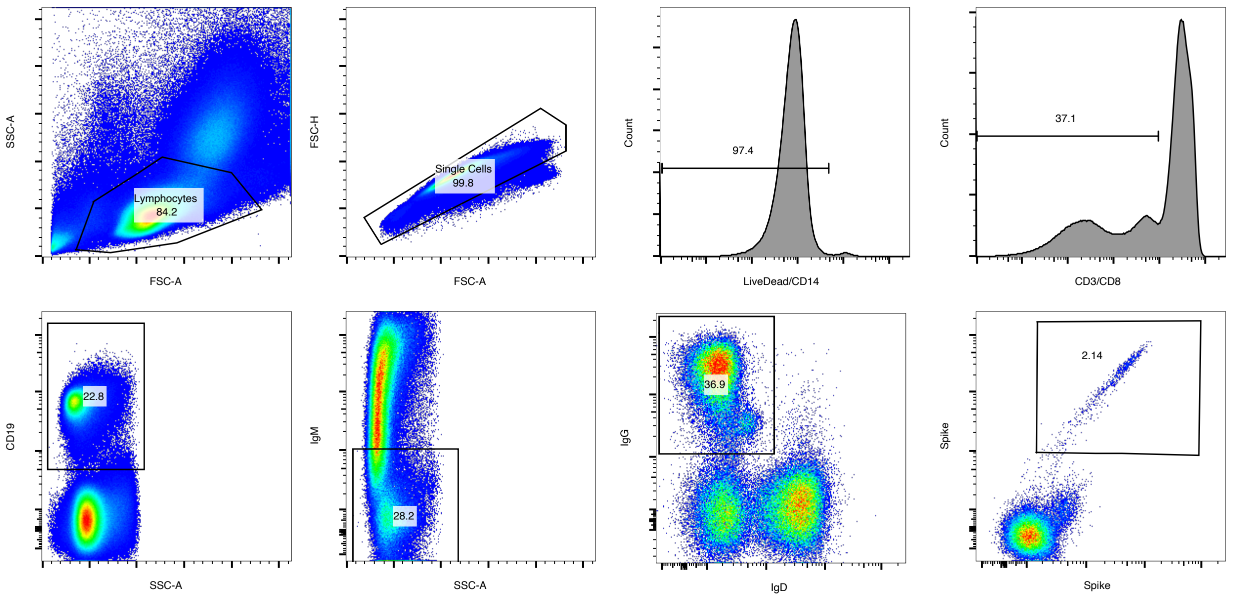

**Supplementary Figure S2: Gene usage for SARS-CoV-2 specific mAbs. A)** Pie charts showing percentage IGHV, IGHD, IGHJ, IGLV/IGLK and IGKJ/IGLJ usage for Spike reactive monoclonal antibodies. **B)** Sankey diagram showing the pairing between VH and VK or VL germline genes for SARS-CoV-2 mAbs isolated. **C)** Bar graph showing the mean VH germline gene percentage usage in SARS-CoV-2 specific mAbs (blue) compared to a representative naïve repertoire (red).[1] Error bars represent the standard deviation between donors used in the analysis (n = 3 for SARS-CoV-2 and n=10 for naïve repertoire). **D)** Single example of clonal expansion observed from P008.

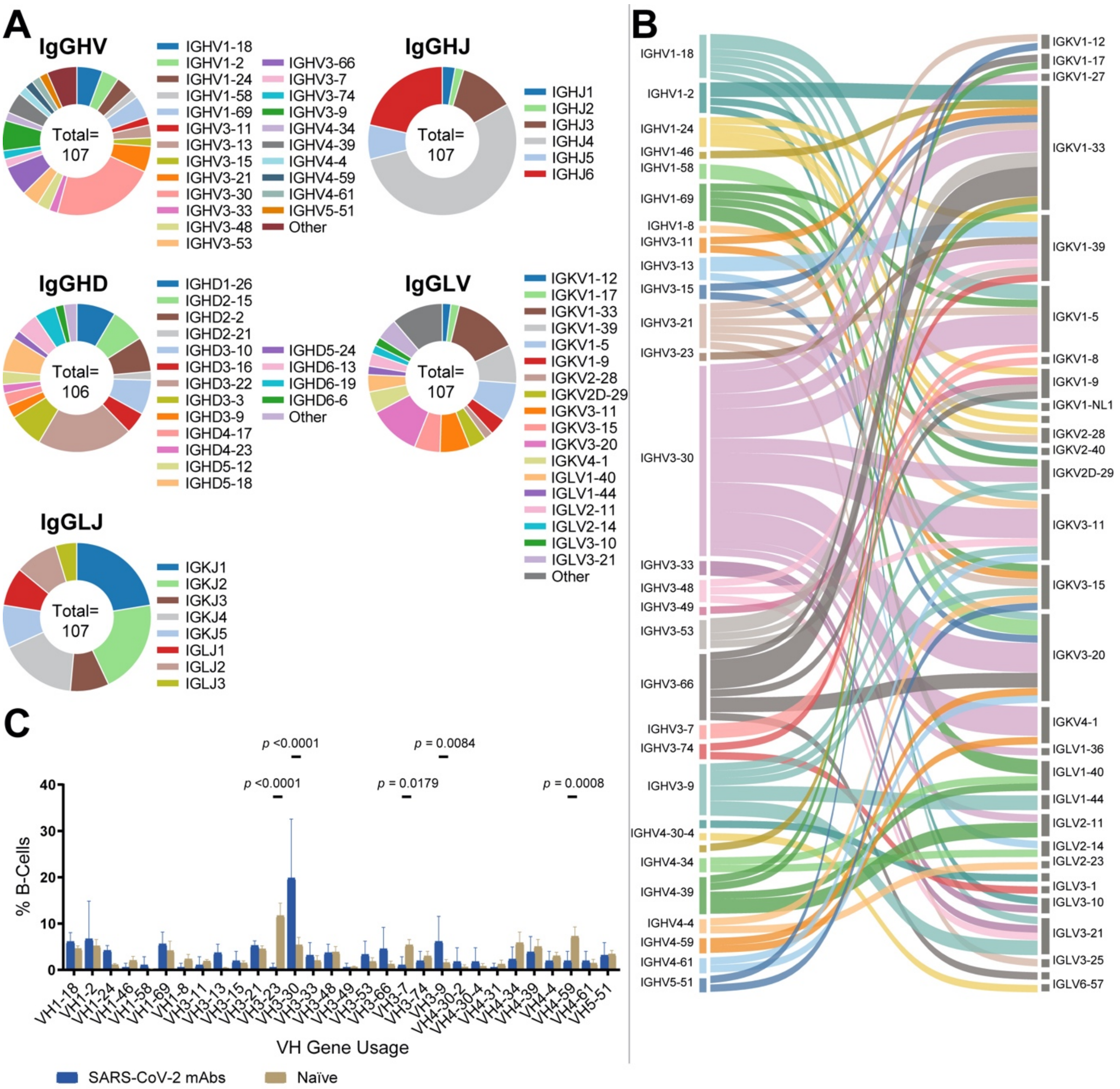


**D**

| Name | VH | JH | VL | JL | Specificity | Neutralization | CDRH3 | CDRL3 | Heavy Chain Sequence Indentity | Light Chain Sequence Indentity |
|---|---|---|---|---|---|---|---|---|---|---|
| P008_036 | IGHV3-30 | IGHJ4 | IGKV2D-29 | IGKJ5 | RBD | Yes | 16 | 9 | 100 | 99.7 |
| P008_044 | IGHV3-30 | IGHJ4 | IGKV2D-29 | IGKJ5 | RBD | Yes | 16 | 9 | | |

**Supplementary Figure S3: Comparison of neutralization of mAbs against SARS-CoV-2 pseudovirus and infectious virus. A)** Correlation of mAb neutralization $IC_{50}$ against infectious virus (y-axis) and pseudotyped virus (x-axis) (Spearman correlation, *r*. A linear regression was used to calculate the goodness of fit, $r^2$). **B)** Example neutralization curves against pseudovirus. **C)** Group 6 nAbs show low neutralization plateaus against pseudovirus. **D)** Unlike the majority of other nAbs, Group 6 nAbs show 5-10 fold higher neutralization potency against infectious virus and reach neutralization plateaus of >95%. Binding of **E)** NTD and RBD nAbs and **F)** S2 non-neutralizing Abs to SARS-CoV-2 and SARS-CoV Spike proteins expressed on the surface of HEK 293T cells measured by flow cytometry. Binding is reported as the % PE-positive cells. Antibodies are colour coded based on their competition group (Group 1 (orange), Group 2 (turquoise), Group 3 (purple), Group 4 (pink), Group 5 (brown), Group 6 (grey), Group 7 (yellow)).

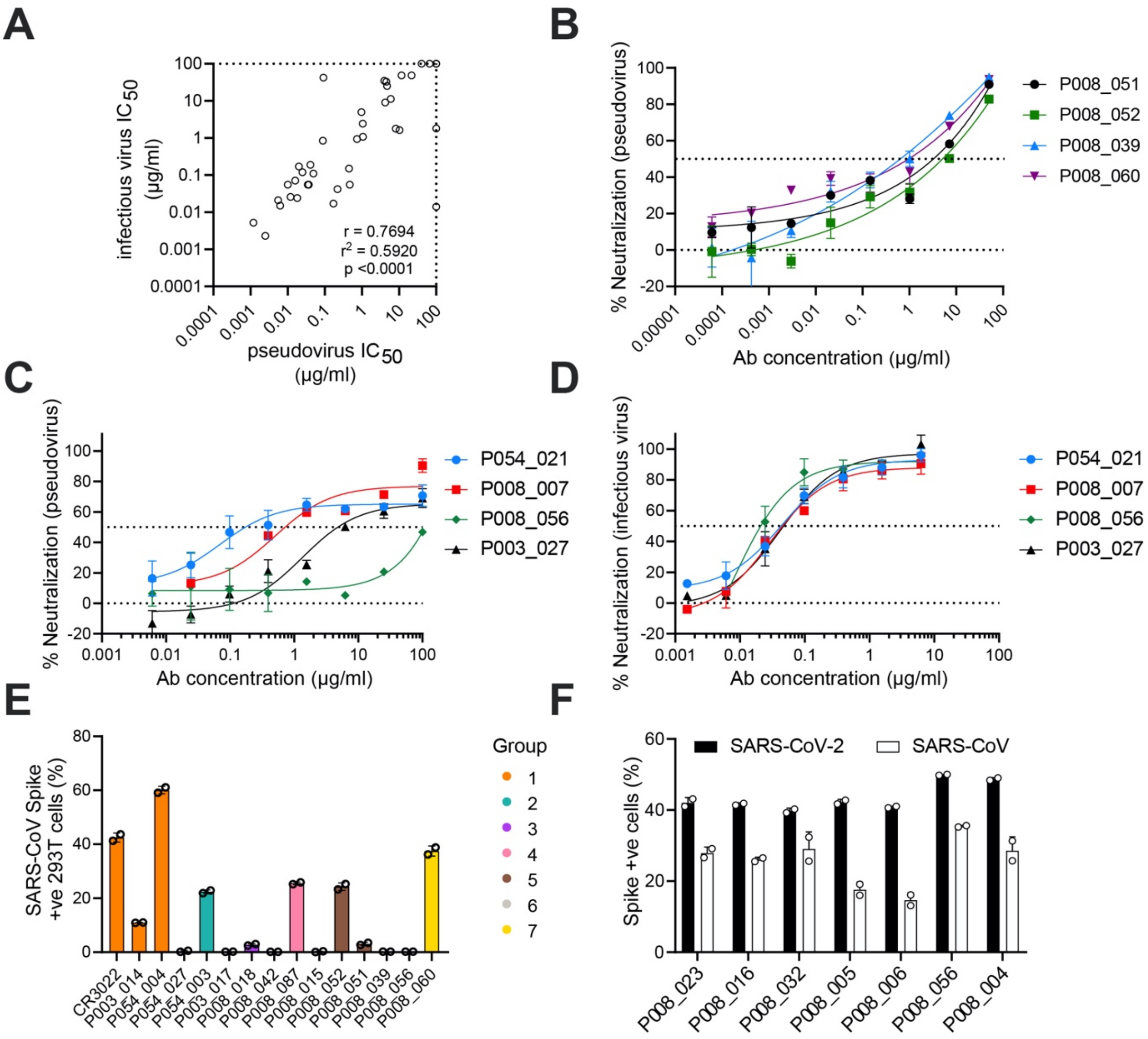
A
infectious virus IC50 (µg/ml)
pseudovirus IC50 (µg/ml)
r = 0.7694
r2 = 0.5920
p <0.0001
B
% Neutralization (pseudovirus)
Ab concentration (µg/ml)
P008_051
P008_052
P008_039
P008_060
C
% Neutralization (pseudovirus)
Ab concentration (µg/ml)
P054_021
P008_007
P008_056
P003_027
D
% Neutralization (infectious virus)
Ab concentration (µg/ml)
P054_021
P008_007
P008_056
P003_027
E
SARS-CoV Spike +ve 293T cells (%)
Group
CR3022
P003_014
P054_004
P054_027
P054_003
P003_017
P008_018
P008_042
P008_087
P008_015
P008_052
P008_051
P008_039
P008_056
P008_060
F
Spike +ve cells (%)
SARS-CoV-2
SARS-CoV
P008_023
P008_016
P008_032
P008_005
P008_006
P008_056
P008_004

**Supplementary Figure S4: Susceptibility of B.1.1.7 and variants to neutralization by neutralizing antibodies.** Neutralization of mAbs and plasma were tested against pseudoviruses expressing variant Spikes including D614G, N501Y and D614G ΔH69/V70 mutations and the B.1.1.7 variant. **A)** Fold change in neutralization of D614G and N501Y mutants compared to wild-type Spike. **B)** Fold change in neutralization of D614G ΔH69/V70 mutant and B.1.1.7 variant compared D614G Spike. A fold change >100 is placed on the dotted line. **C)** Neutralization curves for all mAbs tested. Graphs are arranged by competition group.

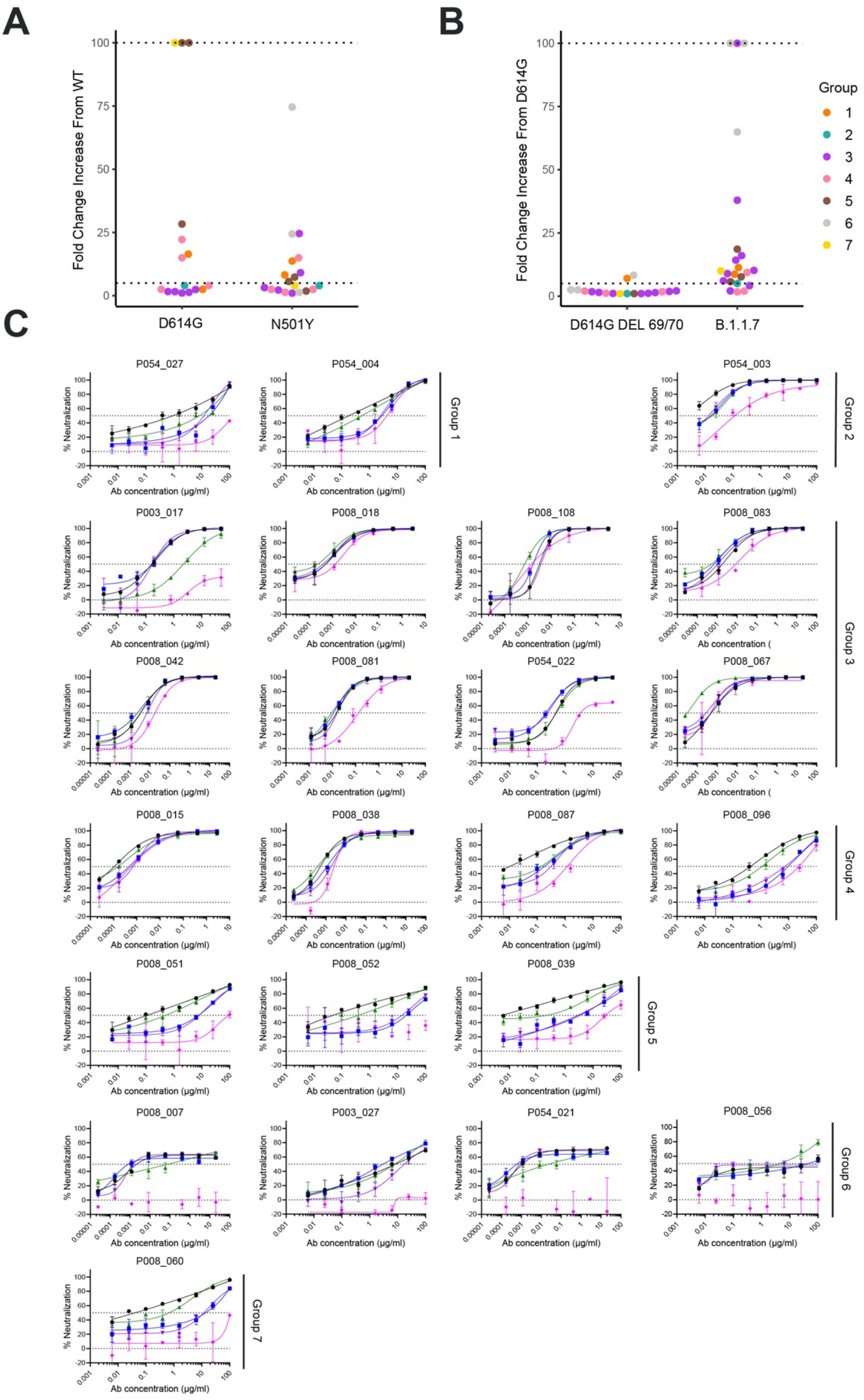
A
B
C
Fold Change Increase From WT
Fold Change Increase From D614G
D614G
N501Y
D614G DEL 69/70
B.1.1.7
Group
1
2
3
4
5
6
7
P054_027
P054_004
P054_003
P003_017
P008_018
P008_108
P008_083
P008_042
P008_081
P054_022
P008_067
P008_015
P008_038
P008_087
P008_096
P008_051
P008_052
P008_039
P008_007
P003_027
P054_021
P008_056
P008_060
% Neutralization
Ab concentration (µg/ml)
Group 1
Group 2
Group 3
Group 4
Group 5
Group 6
Group 7

**Supplementary Table S1: Gene usage, binding characteristics and neutralization properties of SARS-CoV-2 reactive mAbs expressed and purified on large scale.** Antibodies are grouped by donor. Binding competition group for 27 mAbs is listed (see Figure 4A). $EC_{50}$ was measured against Spike, RBD and NTD. For non-S1 binding mAbs, binding to S2 at 25 µg/mL was measured and + indicates binding. Neutralization $ID_{50}$ was measured against infectious virus (with Vero-E6 target cells) and SARS-CoV-2 and SARS-CoV pseudovriuses (with HeLa-ACE2 target cells). The reported $IC_{50}$ values are an average of three independent experiments. %ACE2 competition describes the ability of mAbs to prevent Spike binding to HeLa-ACE2 cells as measured by flow cytometry. The germline VH and VL is reported for each mAb.

| mAb name | Specificty | Competition group | S EC50 | RBD EC50 | NTD EC50 | S2 ELISA | SARS-CoV-2 Infectious Virus IC50 | SARS-CoV-2 pseudovirus IC50 | SARS-CoV pseudovirus IC50 | % ACE2 comp | VH | VL |
|---|---|---|---|---|---|---|---|---|---|---|---|---|
| P003_014 | RBD | 1 | 0.008 | 0.009 | n.d. | n.d. | >100 | >100 | >50 | 86.3 | IGHV3-13 | IGKV1-39 |
| P003_017 | RBD | 3 | 0.006 | 0.007 | n.d. | n.d. | 1.08 | 1.07 | >100 | 96.5 | IGHV3-66 | IGKV1-9 |
| P003_027 | NTD | 6 | 0.004 | >10 | 0.006 | n.d. | 1.64 | 10.30 | >50 | 47.4 | IGHV3-48 | IGKV1-39 |
| P003_016 | NTD | n.d. | 0.130 | >20 | 2.000 | n.d. | 0.153 | 0.449 | >100 | 25.9 | IGHV3-74 | IGKV1-39 |
| P003_055 | NTD | n.d. | 0.060 | >10 | 0.055 | n.d. | >100 | >100 | >100 | n.d. | IGHV4-59 | IGKV4-1 |
| P003_056 | NON-S1 | n.d. | 0.050 | >10 | >10 | + | >100 | >100 | >100 | 36.7 | IGHV3-30 | IGKV4-1 |
| P008_003 | RBD | 3 | 0.076 | 0.083 | n.d. | n.d. | 0.017 | 0.173 | >50 | 99.7 | IGHV4-34 | IGLV1-40 |
| P008_018 | RBD | 3 | 0.071 | 0.082 | n.d. | n.d. | 0.055 | 0.037 | >50 | 99.9 | IGHV3-66 | IGKV3-20 |
| P008_042 | RBD | 3 | 0.128 | 0.173 | n.d. | n.d. | 0.024 | 0.018 | >50 | 99.9 | IGHV3-66 | IGKV3-20 |
| P008_047 | RBD | 3 | 0.173 | 0.233 | n.d. | n.d. | 0.189 | 0.041 | >50 | 99.7 | IGHV3-53 | IGKV1-33 |
| P008_067 | RBD | 3 | 0.164 | 0.186 | >20 | n.d. | 0.170 | 0.020 | >50 | 99.1 | IGHV3-66 | IGKV1-17 |
| P008_081 | RBD | 3 | 0.038 | 0.039 | n.d. | n.d. | 0.116 | 0.025 | >50 | 99.5 | IGHV1-58 | IGKV3-20 |
| P008_083 | RBD | 3 | 0.020 | 0.119 | n.d. | n.d. | 0.026 | 0.012 | >50 | 99.9 | IGHV3-66 | IGKV1-33 |
| P008_090 | RBD | 3 | 0.018 | 0.026 | n.d. | n.d. | 0.071 | 0.016 | >50 | 99.9 | IGHV3-53 | IGKV1-33 |
| P008_108 | RBD | 3 | 0.009 | 0.014 | n.d. | n.d. | 0.0023 | 0.0025 | >50 | 100.0 | IGHV3-11 | IGKV1-33 |
| P008_015 | RBD | 4 | 0.034 | 0.036 | n.d. | n.d. | 0.055 | 0.035 | >50 | 80.2 | IGHV3-9 | IGLV3-21 |
| P008_038 | RBD | 4 | 0.058 | 0.093 | >10 | n.d. | 0.110 | 0.049 | >50 | 42.3 | IGHV1-69 | IGLV1-40 |
| P008_087 | RBD | 4 | 0.021 | 0.056 | n.d. | n.d. | 4.96 | 0.969 | 10.46 | 48.5 | IGHV4-31 | IGKV1-33 |
| P008_096 | RBD | 4 | 0.030 | 0.055 | n.d. | n.d. | 9.09 | 4.26 | >50 | 50.6 | IGHV4-39 | IGKV1-33 |
| P008_057 | RBD | n.d. | 0.032 | 0.060 | >20 | n.d. | 0.015 | 0.006 | >100 | 99.8 | IGHV3-53 | IGKV1-9 |
| P008_070 | RBD | n.d. | 0.034 | 0.108 | n.d. | n.d. | 34.95 | 4.01 | >100 | 47.3 | IGHV3-30 | IGKV1-33 |
| P008_076 | RBD | n.d. | 0.035 | 0.038 | n.d. | n.d. | 0.847 | 0.089 | 0.4648 | 99.0 | IGHV3-13 | IGLV2-14 |
| P008_086 | RBD | n.d. | 0.034 | 0.033 | >20 | n.d. | 0.005 | 0.0012 | >100 | 99.3 | IGHV1-58 | IGKV3-20 |
| P008_100 | RBD | n.d. | 0.033 | 0.036 | >10 | n.d. | 11.30 | 5.98 | >100 | 38.4 | IGHV3-30 | IGKV3-20 |
| P008_103 | RBD | n.d. | 0.066 | 0.101 | n.d. | n.d. | 0.021 | 0.006 | >100 | 99.8 | IGHV3-66 | IGKV1-33 |
| P008_039 | NTD | 5 | 0.101 | >10 | 0.024 | n.d. | 25.28 | 4.78 | >50 | 91.2 | IGHV3-21 | IGKV1-12 |
| P008_051 | NTD | 5 | 0.091 | >10 | 0.056 | n.d. | 48.82 | 21.66 | >50 | 74.5 | IGHV4-61 | IGKV3-11 |
| P008_052 | NTD | 5 | 0.085 | >10 | 0.063 | n.d. | 48.65 | 11.65 | 34.64 | 80.2 | IGHV4-39 | IGLV2-11 |
| P008_007 | NTD | 6 | 0.070 | >10 | 0.171 | n.d. | 0.055 | 0.467 | >100 | 66.3 | IGHV3-48 | IGLV3-21 |
| P008_056 | NTD | 6 | 0.060 | >10 | 0.049 | n.d. | 0.014 | >100 | >50 | 36.8 | IGHV3-21 | IGKV1-33 |
| P008_001 | NTD | n.d. | 0.816 | >20 | 1.200 | n.d. | >100 | >100 | >100 | 37.3 | IGHV3-49 | IGKV1-9 |
| P008_002 | NTD | n.d. | 0.008 | >20 | 0.015 | n.d. | >100 | >100 | >100 | 24.2 | IGHV3-9 | IGLV3-21 |
| P008_014 | NTD | n.d. | 0.066 | >20 | 0.021 | n.d. | 1.81 | >100 | >100 | 43.0 | IGHV3-15 | IGKV1-33 |
| P008_099 | NTD | n.d. | 0.022 | >20 | 0.150 | n.d. | >100 | >100 | >100 | 46.8 | IGHV3-23 | IGKV1-39 |
| P008_060 | NON-S1 | 7 | 0.031 | >10 | >10 | n.d. | 32.90 | 4.52 | 0.6434 | 50.7 | IGHV3-30 | IGKV3-20 |
| P008_004 | NON-S1 | n.d. | 0.045 | >10 | >10 | + | >100 | >100 | >100 | 61.4 | IGHV3-7 | IGKV1-5 |
| P008_005 | NON-S1 | n.d. | 0.200 | >10 | >10 | + | >100 | >100 | >100 | 22.8 | IGHV3-30 | IGKV1-5 |
| P008_006 | NON-S1 | n.d. | 0.100 | >10 | >10 | + | >100 | >100 | >100 | 16.0 | IGHV3-30 | IGKV1-5 |
| P008_016 | NON-S1 | n.d. | 0.108 | >10 | >10 | + | >100 | >100 | >100 | 39.9 | IGHV4-59 | IGKV3-20 |
| P008_023 | NON-S1 | n.d. | 0.400 | >10 | >10 | + | >100 | >100 | >100 | 22.7 | IGHV3-11 | IGKV3-15 |
| P008_032 | NON-S1 | n.d. | 0.060 | >10 | >10 | + | >100 | >100 | >100 | 23.3 | IGHV3-30 | IGKV3-11 |
| P054_004 | RBD | 1 | 0.036 | 0.051 | n.d. | n.d. | 42.23 | 0.092 | 5.9585 | 95.2 | IGHV4-34 | IGLV2-14 |
| P054_027 | RBD | 1 | 0.008 | 0.013 | n.d. | n.d. | >100 | 40.32 | >50 | 92.9 | IGHV4-30-4 | IGLV6-57 |
| P054_003 | RBD | 2 | 0.176 | 0.200 | n.d. | n.d. | 1.79 | 8.20 | >100 | 98.5 | IGHV1-2 | IGKV1-33 |
| P054_022 | RBD | 3 | 0.174 | 0.194 | n.d. | n.d. | 2.440 | 1.048 | >50 | 98.5 | IGHV4-30-2 | IGLV2-8 |
| P054_036 | RBD | n.d. | 0.013 | 0.029 | >10 | n.d. | 0.055 | 0.010 | >100 | 99.5 | IGHV3-53 | IGKV1-39 |
| P054_050 | RBD | n.d. | 0.027 | 0.047 | n.d. | n.d. | >100 | 66.60 | >100 | 60.1 | IGHV3-13 | IGKV1-39 |
| P054_021 | NTD | 6 | 0.026 | >10 | 0.064 | n.d. | 0.042 | 0.218 | >100 | 51.0 | IGHV1-24 | IGKV2-24 |
| P054_044 | NTD | n.d. | 0.031 | >20 | 4.000 | n.d. | 0.943 | 0.740 | >100 | 46.7 | IGHV1-69 | IGKV1-5 |
